# Anchoring of perforin-2 via the transmembrane domain is required for endocytic escape in cross-presenting dendritic cells

**DOI:** 10.64898/2025.12.03.692202

**Authors:** Marco Laub, Ritika Chatterjee, Patrycja Kozik

## Abstract

Perforin-2 is a pore-forming protein localised to the endocytic compartments of dendritic cells and macrophages. It is reported to perform two distinct functions during immune responses: attacking intravacuolar pathogens, and forming pores in endocytic compartments to enable cytosolic delivery of antigens during cross-presentation. The molecular mechanisms that regulate perforin-2 remain unknown. Here, we set out to address how cross-presenting dendritic cells control pore formation in phagosomes while maintaining the integrity of their endocytic compartments. We demonstrate that perforin-2 undergoes extensive proteolytic processing involving multiple endocytic proteases. Although the transmembrane anchor has been proposed to protect host membranes by orienting pores towards bacterial targets, we find that endocytic escape is mediated by full-length, membrane-anchored perforin-2 rather than by the proteolytically released ectodomain. Moreover, we show that perforin-2-mediated antigen translocation does not require low pH, explaining how perforin-2 mediated antigen import into the cytosol can occur in cross-presenting dendritic cells which do not acidify their phagosomes. Our findings point at a critical role of the transmembrane anchor in perforin-2 in dendritic cells, and reveal that pore formation occurs with striking spatiotemporal precision to safeguard the integrity of endocytic compartments.

## Introduction

The mammalian immune system employs a family of membrane attack complex and perforin (MACPF) domain-containing pore-forming proteins in innate and adaptive immunity. The family includes C6-C9 subunits of the complement membrane attack complex as well as two perforins. Complement components are produced primarily by hepatocytes and secreted into the serum, where they can form lytic pores on the surface of Gram-negative bacteria, enveloped viruses, parasites, and host cells. Perforin-1 (*Prf1*) is expressed in cytotoxic T cells and natural killer cells. Upon activation, perforin-1-containing cytotoxic granules are secreted into the immune synapse between the killer cell and the target. Perforin-1 pores can be lytic at high concentration, but their primary role is to deliver apoptosis-inducing granzymes into the cytosol of the target cell (reviewed in (Krawczyk *et al*, 2020)). Perforin-2 (*Mpeg1*) is the evolutionarily most ancient member of the mammalian MACPF family (McCormack & Podack, 2015), yet it remains the least well characterised. Perforin-2 is highly expressed by antigen presenting cells such as dendritic cells and macrophages as well as by neutrophils. Initially, it has been proposed to target intravacuolar bacteria (Fields *et al*, 2013; McCormack *et al*, 2015) , but its role in anti-bacterial immunity has been questioned in later work (Ebrahimnezhaddarzi *et al*, 2022). More recently, we demonstrated that perforin-2 can form pores in the membranes of endocytic compartments to facilitate the release of internalised antigens into the cytosol, where they are processed by the proteasomes (Rodríguez-Silvestre *et al*, 2023). Proteasome-derived peptides are then presented on MHC class I molecules, enabling APCs to activate antigen-specific T cells even when they do not express the antigens themselves, in a pathway referred to as cross-presentation.

The mechanism of pore formation by MACPF-containing proteins is broadly conserved and shared with a related family of cholesterol-dependent cytolysins (CDCs) present in bacteria (Dunstone & Tweten, 2012). The key steps of this process include 1) association with the membrane, 2) oligomerisation into a pre-pore structure and 3) transition from the pre-pore to pore conformation through unwinding of the α-helical transmembrane hairpins (TMH1 and TMH2). Following the conformational change, each TMH contributes two β-sheets to the resulting β-barrel pore (Serna *et al*, 2016; Law *et al*, 2010; Ivanova *et al*, 2022; Pang *et al*, 2019; Ni *et al*, 2020). Distinct from other proteins in the MACPF/CDC superfamily, perforin-2 has an additional transmembrane domain (TMD) which does not form part of the pore (Pang *et al*, 2019; Ni *et al*, 2020) . As a consequence of this membrane anchoring, in the oligomeric pre-pore state of perforin-2, the pore-forming TMH helices point away from the membrane (Figure 1A). Thus, it has been initially proposed that the TMD serves to protect endogenous membranes by orienting the pores towards the lumen of a pathogen-containing vacuole (Pang *et al*, 2019). However, perforin-2’s function in endocytic escape of antigens in cross-presenting DCs suggests that pore formation in endogenous membranes is an important function of perforin-2, rather than an unwanted effect (Rodríguez-Silvestre *et al*, 2023).

**Figure 1.**
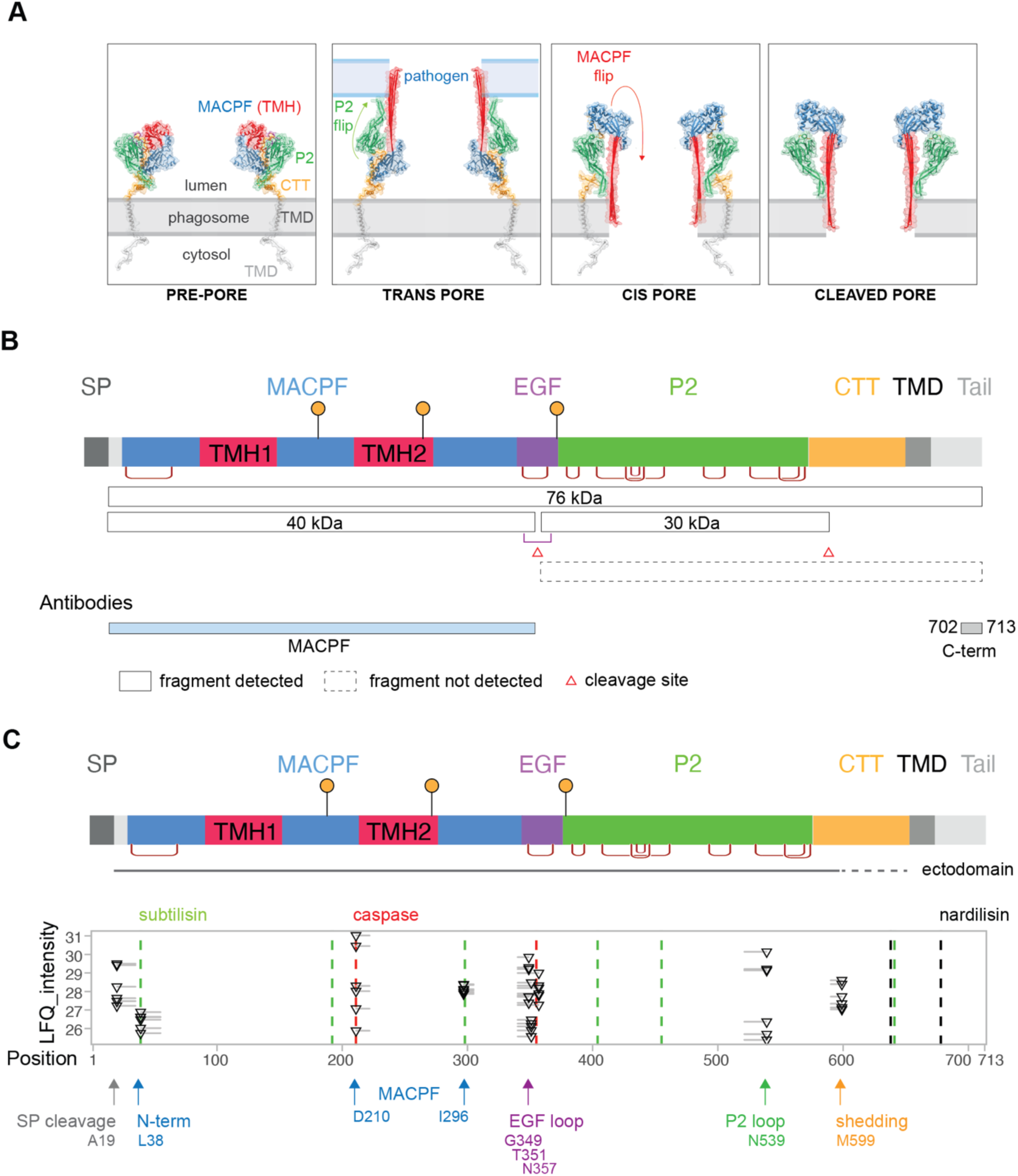
Mapping of perforin-2 cleavage sites by proteomics. **A** Model of the pre-, trans-, cis- and cleaved pores of perforin-2. **B** Schematic diagram illustrating hypothetical (dashed lines) and previously observed (solid lines) perforin-2 fragments. Epitope for custom perforin-2 antibody is shown in colour. **C** Non-tryptic peptides identified by mass-spectrometry of total cell lysates of MutuDCs. The triangles indicate non-tryptic cleavage sites. Vertical dotted lines indicate cleavage sites predicted by ELM database.

If TMHs face into the organellar lumen, how are perforin-2 pores inserted into the membranes during cross-presentation? One solution to this topological problem is to proteolytically release perforin-2’s ectodomain from the membrane. Notably, the structures of perforin-2 pores have been obtained using the ectodomain only, suggesting, at least *in vitro*, perforin-2 can undergo pre-pore to pore transition without the TMD (Pang *et al*, 2019; Ni *et al*, 2020; Yu *et al*, 2022). Indeed, in the cDC1-like cell line, MutuDCs (Marraco *et al*, 2012), the majority of perforin-2 has lost the TMD (Rodríguez-Silvestre *et al*, 2023; Ebrahimnezhaddarzi *et al*, 2022). Such proteolytic release would enable rotation of the TMHs towards endogenous membranes. However, in the absence of the TMD, perforin-2 binds membranes via the tip of the P2 domain, which in the pre-pore points in the opposite direction to the TMHs (see Figure 1). Thus, the proteolytic release alone does not explain how the pores can form on the same membrane on which the pre-pore sits.

Interestingly, two models have been proposed for the pre-pore to pore conformational transition of perforin-2. The model where the TMHs unfurl away from the membrane and insert into an opposing bilayer is referred to as the trans model (Figure 1A). Whether formation of the trans pores is preceded by the rotation of the P2 domain has not been demonstrated, but it appears likely, as it would facilitate binding (and pore insertion) to the target membrane (Pang *et al*, 2019; Ni *et al*, 2020). This mode of pore formation would allow for targeting intravacuolar pathogens, but, as discussed above, is unlikely to facilitate the release of endocytic cargo. In the alternative cis-model, the P2 domain remains bound to the membrane, but the MACPF domain rotates by 180 degrees prior to unfurling (Figure 1A) (Jiao *et al*, 2022; Yu *et al*, 2022). This allows for the insertion of pores into the membrane on which the pre-pore sits. The cis-model could be employed during pore formation on bacterial membranes, but this would require proteolytic release of the ectodomain to enable pre-pores to bind the pathogen surface. Importantly, this model would also enable the formation of pores in the membranes of endocytic compartments. Theoretically, the TMD would not need to be removed to enable endocytic escape via cis pores, but whether full length perforin-2 can oligomerise and make pores has not been demonstrated.

Here, we asked whether ectodomain shedding is required for perforin-2 mediated endocytic escape. To this end, we first mapped the cleavage sites involved in perforin-2 maturation and identified the proteases involved. This enabled us to inhibit proteolytic maturation of perforin-2. Surprisingly, we found that shedding was not required for pore formation. Instead, we demonstrate that membrane anchoring of perforin-2 facilitates formation of endocytic escape pores, and mutations in the CTT domain, which links the ectodomain with the TMD, interfere with escape. Together, these data point at a central role of the TMD anchor during perforin-2 mediated antigen escape and indicate that ectodomain shedding serves as a mechanism to inactivate perforin-2 in dendritic cells.

## Results

### Mapping of perforin-2 cleavage sites by proteomics

The perforin-2 ectodomain has a predicted molecular weight (mw) of approximately 76 kDa and is composed of the pore-forming MACPF domain, an epidermal growth factor (EGF) linker, a membrane binding P2 domain, and an unstructured CTT domain that connects it to the TMD (Figure 1B). At steady state, in addition to full-length perforin-2, two fragments have been robustly detected in murine DCs: a ∼40 kDa MACPF domain (consistent with the predicted mw of 35.5 kDa plus two N-linked glycans), and a ∼30 kDa P2 (theoretical mw 22-30 kDa, depending on the precise location of the shedding site, plus one N-linked glycan) (Figure 1B) (Rodríguez-Silvestre *et al*, 2023; Ebrahimnezhaddarzi *et al*, 2022). Because the antibody against the C-terminal tail does not recognise the ∼30 kDa P2 fragment, these observations suggest that two cleavage sites are present in perforin-2: one in the EGF domain resulting in two fragments linked by a disulfide bond, and one responsible for ectodomain shedding (Figure 1B).

To define the proteolytic steps involved in perforin-2 maturation, we first mapped candidate cleavage sites using previously published mass spectrometry data from whole cell lysates of MutuDC (Rodríguez-Silvestre *et al*, 2023). Figure 1C shows the positions of non-tryptic peptides - *i.e.* peptides likely generated in cells rather than produced by trypsin digestion of the lysates - identified in each of six independent samples. As a complementary approach, we used the Eukaryotic Linear Motif (ELM) database (Kumar *et al*, 2023) to identify known protease recognition motifs in the mouse perforin-2 sequence (indicated with dashed lines) (Figure 1C, Table S1).

In addition to the previously reported promiscuous cleavage in the EGF loop (Rodríguez-Silvestre *et al*, 2023; Ebrahimnezhaddarzi *et al*, 2022), we observed five putative cleavage events: (1) a site at position L38^, which would remove a short N-terminal fragment; (2-3) two sites within the MACPF domain (D210^ and I296^); and (4) one site in the P2 loop (N539^); (5) a candidate shedding site in the CTT at M599^ (cleavage at position A19^ corresponds to signal peptide removal). Three of the five sites in the ectodomain (N-term, EGF and the P2 loop cleavages) are between cysteine residues linked by disulfides bond, and are therefore unlikely to result in major structural rearrangements in non-reducing conditions (Figure 1C). We hypothesize that the two cleavages in the MACPF domain (D210^ and I296^) are involved in the disassembly of perforin-2, as they would compromise the integrity of the pore-forming domain. In line with this hypothesis, cleavage at D210^ corresponds to a putative caspase site and is therefore likely to occur following pore formation, when the β-barrel is accessible from the cytosol. The candidate shedding site is in the unstructured region of the CTT, 55 residues away from the transmembrane domain. In summary, the proteomics-based analysis uncovers a previously unappreciated level of proteolytic remodelling of perforin-2, which includes, but is not limited to, ectodomain shedding.

### P2 loop is cleaved by Asparagine Endopeptidase (AEP)

To validate the cleavage sites identified by proteomics, we raised antibodies against the N-terminal end of perforin-2 (residues D21-C34) and the P2 loop (residues N539-C554) (Figure 2A). The N-terminal antibody detected full-length perforin-2, but it did not recognise the 40 kDa MACPF fragment revealed by a commercial antibody with an unknown binding site (Figure 2B-C). Thus, these data confirm that the N-terminus is cleaved at an early stage of perforin-2 maturation, and suggest that this occurs prior to or concurrently with cleavage in the EGF domain.

**Figure 2.**
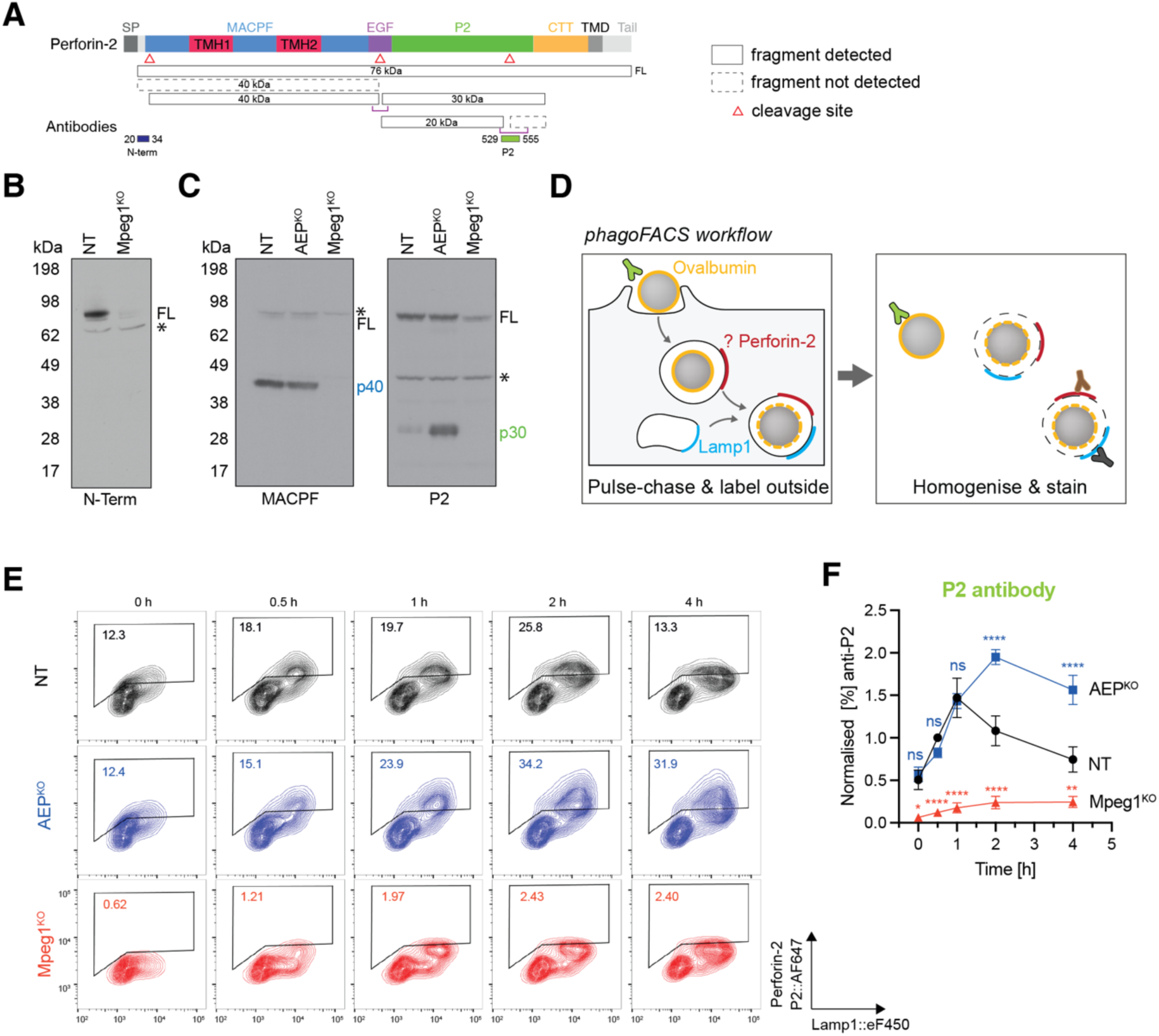
Perforin-2 is cleaved by AEP in the EGF and P2 domains. **A** Diagram illustrating the AEP cleavage sites and the αP2 antibody epitope (black) in the pre-pore (PDB 6SB3) and pore (PDB 6SB5) structures of the perforin-2 ectodomain. **B** Representative immunoblot analysis under reducing conditions of lysates of NT, and Mpeg1^KO^ MutuDCs using the αN-term antibodies. * indicates non-specific bands. The corresponding Ponceau blot is shown in Figure S1. **C** Representative immunoblot analysis under reducing conditions of lysates of NT, AEP^KO^ and Mpeg1^KO^ MutuDCs using the αMACPF and αP2 antibodies. The corresponding ponceau blot is shown in Figure S1. **D** Diagram illustrating the phagoFACS experimental setup: NT, AEP^KO^ and Mpeg1^KO^ MutuDCs were pulsed with OVA beads and chased for the indicated time. Isolated phagosomes were stained with αP2 and αLamp1 antibodies. **F** Quantification αP2+ phagosomes from phagoFACS experiment as gated in (**E**), normalised to control at 1 hour. Mean and SEM. For all data, n = 3 independent experiments.

The antibody against the P2 loop detected full-length perforin-2 as well as a faint band corresponding to the P2 domain at 30 kDa (Figure 2C). We have previously demonstrated that the EGF-linker is cleaved by asparagine endopeptidase (AEP). In the absence of AEP, the EGF-linker is still cleaved, but the cleavage results in different tryptic fragments, suggesting that other proteases can compensate for AEP activity (Rodríguez-Silvestre *et al*, 2023). Data from AEP knockout cells also suggested that AEP might control an additional cleavage, located in the P2 loop, as the non-tryptic peptide p521-539 was depleted in the AEP^KO^ cells (Rodríguez-Silvestre *et al*, 2023). Indeed, in the absence of AEP, we observed accumulation of the 30 kDa P2 domain (Figure 2C) when probing with the N539-C554 P2 antibody. The cleavage in the P2 domain is corroborated by the previous observation of a faint 20 kDa fragment detected with a commercial αP2 antibody (Rodríguez-Silvestre *et al*, 2023). The immunoblotting data also suggest that cleavage in the P2 loop occurs after (or at the same time as) the EGF loop cleavage, as there is no accumulation of full-length perforin-2 (nor appearance of any other fragments) in the AEP^KO^ cells under reducing conditions (see also Figure S8).

To test whether the P2 loop cleavages occur following recruitment to antigen-containing compartments, we analysed perforin-2 processing in phagosomes. We pulsed the cells with ovalbumin (OVA)-coated beads, and at different time points, labelled outside beads with anti-OVA antibody and lysed the cells. The phagosomes were then isolated and stained with epitope-specific antibodies against perforin-2 and Lamp1 (Figure 2D, S2A). As we reported previously, the signal from the C-term antibody, which detects full-length perforin-2, peaked within 30 min (Figure S2B,C) and was then lost over time, most likely due to removal of the TMD from the membrane following ectodomain shedding. The signal from the P2-loop antibody peaked later, at 2 hours (Figure 2E-F), suggesting the P2 loop is cleaved at slower kinetics compared to shedding. In AEP^KO^ cells, initial recruitment of perforin-2 recognised by the P2-loop antibody follows similar kinetics to control cells, but at later time points, the signal continues to accumulate, consistent with the model where, in the absence of AEP, the P2 loop is no longer cleaved.

Together, these data demonstrate that N-terminal cleavage represents an early maturation event, whereas cleavage within the P2 domain occurs once perforin-2 is recruited to phagosomes and has undergone ectodomain shedding.

### TMD and the cytosolic tail are removed from the membrane by γ-secretase

As we were unable to detect any fragments of cleaved perforin-2 with the C-terminal tail antibody, we predicted that following shedding, the TMD and the cytosolic tail are rapidly removed from the membranes. The best characterised intramembrane protease in the endocytic compartments is y-secretase, which recognises substrates once their luminal domain has been removed (Kopan & Ilagan, 2004). Indeed, when we treated the cells with y-secretase inhibitors (DAPT, LY-411,575, and LY-685,458), we observed accumulation of a 12 kDa fragment detected with the C-term antibody (Figure 3A,B). The size of the fragment is consistent with the predicted shedding site in the CTT domain, at position M599^. We also confirmed these results in MutuDC lines deficient in γ-secretase activity generated by knocking out the auxiliary subunit nicastrin (Ncstn) (Figure 3C). Knockout of the proteolytically active subunit, presenilin (Psen2), or the anterior pharynx defective 1 (Aph1c) did not abolish the y-secretase mediated cleavage, most likely due to the homologous genes Psen1 and Aph1a/Aph1c, respectively (Figure 3C). In cells treated with y-secretase inhibitor DAPT, we also observed accumulation of the C-terminal tail staining on phagosomes, in line with the hypothesis that the TMD and the cytosolic tail are removed from the membrane during phagosome maturation (Figure 3D-E). These data are consistent with the model in which full-length perforin-2 is recruited phagosomes, where the ectodomain is shed into the lumen and the TMD with the cytosolic tail are removed from the membrane by y-secretase.

**Figure 3.**
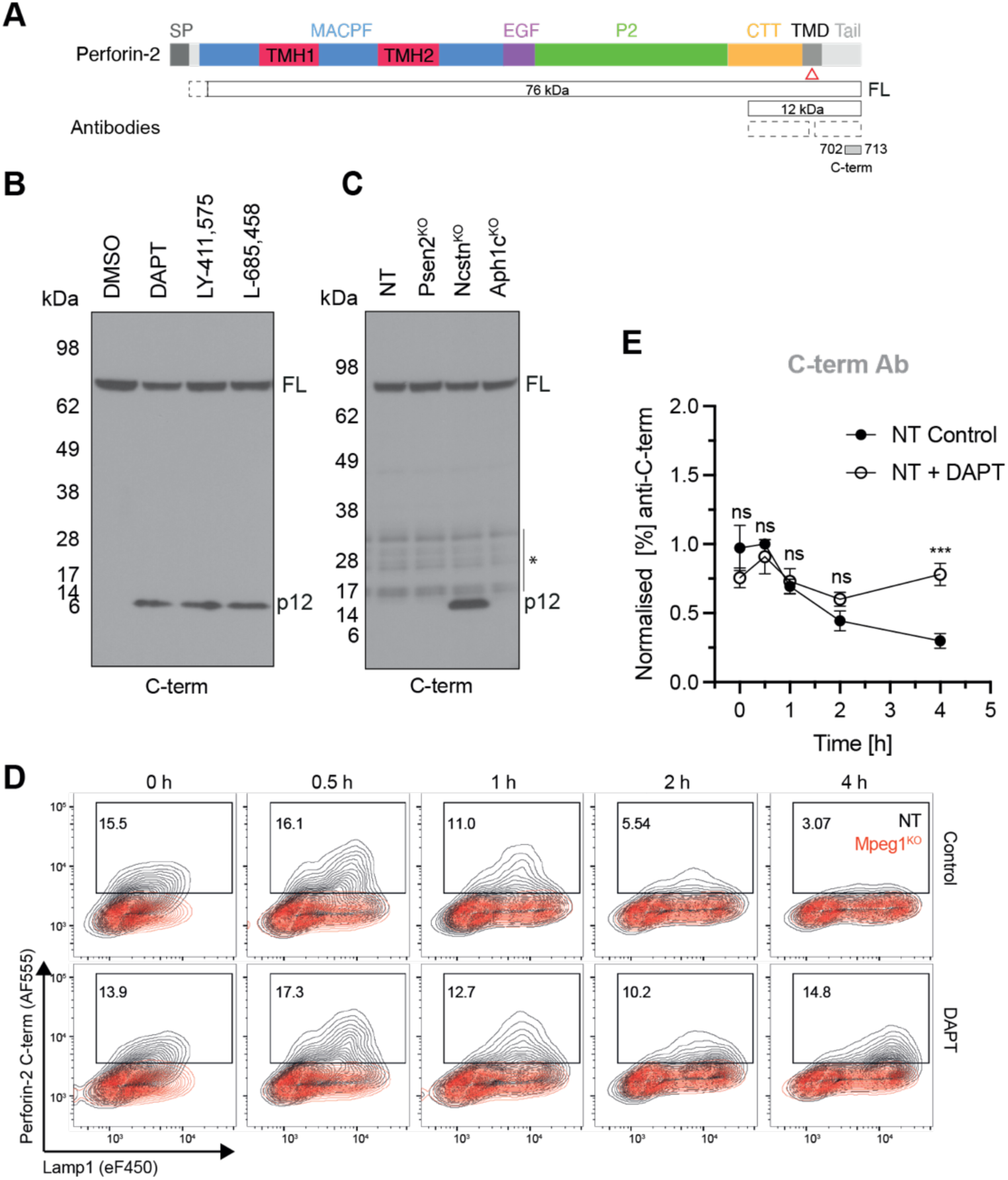
Perforin-2 is cleaved by γ-secretase. **A** Diagram illustrating the approximate cleavage sites (indicated with triangles) of γ-secretase and of the putative sheddase. **B** NT MutuDCs were treated with either DMSO (0.1% v/v), DAPT (10 μM), LY-411,575 (10 μM) or L-685,458 (10 μM) for 8 hours and lysates analysed by immunoblot under non-reducing conditions using the αC-terminal tail antibody. n = 10 independent experiment for DAPT conditions and the data representative of n = 2 independent experiments for LY-411,575 and L-685,458 conditions. The corresponding Ponceau blot is shown in Figure S1. **C** Lysates of NT, Psen2^KO^, Ncstn^KO^ and Aph1c^KO^ MutuDCs were analysed by immunoblot under non-reducing conditions using the αC-terminal tail antibody. Data representative of n = 2 independent experiments. The corresponding Ponceau blot is shown in Figure S1. **D** NT and Mpeg1^KO^ MutuDCs were pulsed with OVA beads and chased in the presence or absence of DAPT (10 μM) for the indicated time. Isolated phagosomes were stained with αC-term tail and αLamp-1 antibodies. **E** Quantification αC-terminal tail + phagosomes as gated in (D) normalised to control at 1 hour. Data are mean and SEM. n = 3 independent experiments.

### Shedding is inhibited by broad cathepsin inhibitor, Z-FA-FMK

In order to identify the mechanism of shedding, we screened small-molecule inhibitors for those that prevent the appearance of the 12 kDa perforin-2 TMD with the cytosolic tail in the presence of DAPT (Figure 1A). If the responsible sheddase were inhibited, the p12 fragment should no longer be generated and therefore would not accumulate upon γ-secretase inhibition. Most known sheddases belong to “a disintegrin and metalloprotease” (ADAM), beta-site APP cleaving enzyme (BACE) or matrix metalloprotease (MMP) families (Lichtenthaler *et al*, 2018). We therefore pre-treated MutuDCs for 2 hours with a panel of inhibitors targeting these enzymes (Figure 4B), followed by a 6-hour incubation in the presence of DAPT. However, none of the classical sheddase inhibitors prevented generation of the p12 fragment (Figure 4C).

**Figure 4.**
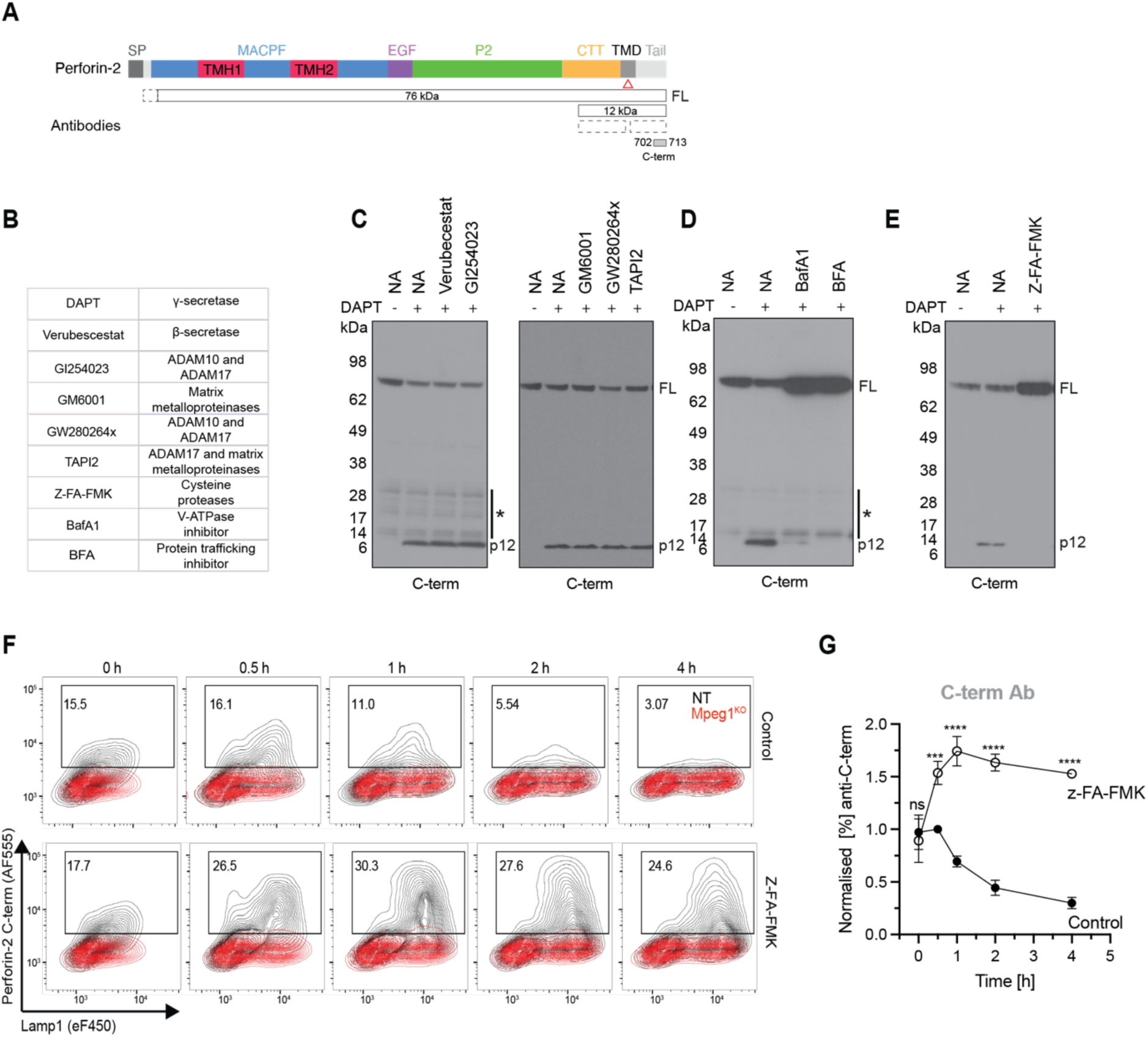
Ectodomain shedding is executed by cathepsins in acidified compartments. **A** Diagram illustrating the approximate cleavage sites of small molecule inhibitors **B** Small molecule inhibitors and their targets **C** NT MutuDCs were either left untreated, treated with DAPT (10 μM) alone for 8 hours or pre-treated with 1 µM Verubecestat or 10 µM GI254023 or 10 µM GM6001, 10 µM GW280264x or 20 µM TAPI2 for 2 hours, followed by a 6-hour treatment with DAPT (10 μM) in the continued presence of respective inhibitors. Lysates were analysed by immunoblot under non-reducing conditions using the αC-terminal tail antibody. The corresponding Ponceau blot is shown in Figure S1. **D** NT MutuDCs were either left untreated, treated with DAPT (10 μM) alone for 8 hours or pre-treated with bafilomycin A1 (0.5 μM) or brefeldin A (5 μM) for 2 hours, followed by a 6-hour treatment with DAPT (10 μM) in the continued presence of bafilomycin A1 or brefeldin A. Lysates were analysed by immunoblot under non-reducing conditions using the αC-terminal tail antibody. The corresponding Ponceau blot is shown in Figure S1. **E** As in A except that NT MutuDCs were pre-treated with Z-FA-FMK (10 μM). The corresponding Ponceau blot is shown in Figure S1, and the data are representative of n =2 independent experiments. **F** NT and Mpeg1^KO^ MutuDCs were pulsed with OVA beads and chased in the presence or absence of Z-FA-FMK (10 μM) for the indicated time. Isolated phagosomes were stained with αC-term tail and αLamp1 antibodies. **G** Quantification αC-terminal tail+ phagosomes as gated in (D) normalised to control at 1 hour. Data are mean and SEM. n = 2 independent experiments.

We then tested whether ectodomain shedding depends on perforin-2 delivery to phagosomes or phagosome acidification. We treated MutuDCs with an inhibitor of ARF GTPases, Brefeldin A, which blocks transport through the secretory pathway as well as with V-ATPase inhibitor, Bafilomycin A1. Both treatments prevented generation of the p12 fragment and resulted in accumulation of full-length perforin-2 (Figure 4D), suggesting that shedding might be mediated by lysosomal proteases. Indeed, we were able to inhibit shedding with Z-FA-FMK, which primarily targets cysteine proteases, including cathepsin B and L (Hara *et al*, 1997; Citarella & Micale, 2020; Lopez-Hernandez *et al*, 2003). In line with these data, Z-FA-FMK resulted in strong accumulation of perforin-2 C-term staining on phagosomes (Figure 4F-G). Together, these results suggest that Z-FA-FMK targets the protease(s) responsible for perforin-2 ectodomain shedding and that shedding occurs once perforin-2-containing phagosomes acidify.

### Ectodomain shedding is not required for endocytic escape

To test whether proteolytic maturation of perforin-2 is required for pore formation in the phagocytic membranes, we employed the saporin-based endocytic escape assay (Figure 5A) (Rodríguez-Silvestre *et al*, 2023). Saporin is a ribosome-inactivating toxin (RIP), which when delivered into the cytosol, leads to translation arrest (Nielsen & Boston, 2001). To monitor endocytic escape from phagosomes, we conjugated saporin to ovalbumin-coated beads using the heterobifunctional cross-linker SPDP, which introduces disulphide bonds. The bond can be reduced through the activity of phagosomal Gamma-interferon–inducible lysosomal thiolreductase (GILT), resulting in the release of saporin into the phagosome lumen (Huotari & Helenius, 2011). To monitor saporin-mediated translation arrest, we labelled nascent polypeptide chains with puromycin (Schmidt *et al*, 2009). In line with previous data (Rodríguez-Silvestre *et al*, 2023), saporin escape from the beads was abolished in Mpeg1^KO^ MutuDCs (Figure 5B, S3A). As reported before, in the saporin-puromycin assay, we observe two populations of cells: cells with efficient translation, and cells where saporin escaped into the cytosol, leading to translation arrest (Figure 5B). This distribution of cells into two populations suggests that, once a small number of ribosomes are inactivated, there is a translational shutdown, possibly due to a ribotoxic stress response (Vind *et al*, 2020; Nielsen & Boston, 2001). To understand how the percentage of in cells in translation arrests correlates to the efficiency of saporin escape, we titrated the amount of saporin on the beads (Figure 5B-D). In cells containing single phagosome, a 2-fold decrease in the amount of saporin on the beads resulted in about 4% decrease in the percentage of cells in translation arrests, demonstrating that the assay is quantitative and that in Mpeg1^KO^ MutuDC antigen escape is at least 30-fold less efficient.

**Figure 5.**
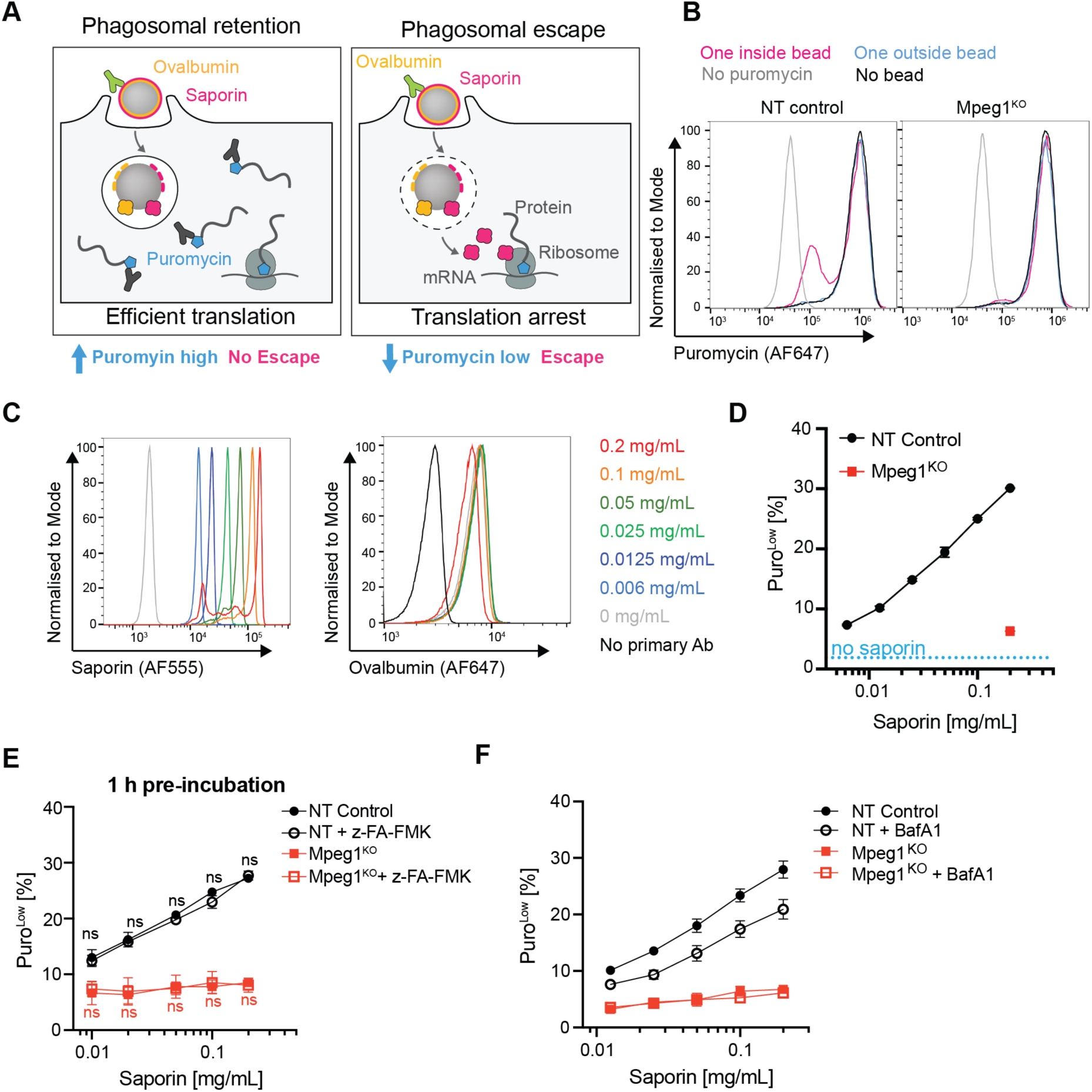
Proteolytic processing of perforin-2 is not required for endocytic escape. **A** Schematic representation of the bead saporin-puromycin assay. **B** NT or Mpeg1^KO^ MutuDCs were incubated with saporin beads for 5 h followed by an incubation with puromycin for 30 min. Cells were then placed on ice and outside beads labelled with an αOvalbumin antibody. After fixation and permeabilisation, levels of puromycin incorporation were detected with an αPuromycin antibody. Data are representative of n =5 independent experiments. **C** Amino beads were coated in ovalbumin followed by conjugation with different concentrations of saporin as indicated. Beads were then stained with αSaporin and αOvalbumin antibodies. **D** NT or Mpeg1^KO^ MutuDCs were incubated with saporin beads for 5 h followed by an incubation with puromycin for 30 min. Cells were then placed on ice and outside beads labelled with an αOvalbumin antibody. After fixation and permeabilisation, levels of puromycin incorporation were detected with an αPuromycin antibody. Quantification of cells in translation arrest (PuroLow). Mean of two technical repeats. **E** NT or Mpeg1^KO^ MutuDCs were incubated with saporin beads for 5 h followed by an incubation with puromycin for 30 min and pre-incubated for 1h with Z-FA-FMK. Cells were then placed on ice and outside beads labelled with an αOvalbumin antibody. After fixation and permeabilisation, levels of puromycin incorporation were detected with an αPuromycin antibody. Quantification of cells in translation arrest (PuroLow) based on the gating shown in (Supp Figure 3B). Data represent mean and SEM of n = 2. **F** NT and Mpeg1^KO^ MutuDCs were assayed in the bead saporin-puromycin assay in the presence or absence of 0.5 µM bafilomycin A1. Quantification of cells in translation arrest (PuroLow) based on the gating shown in (Supp Figure 3C). Data represent mean and SEM of three independent experiments with BafA1-treated Mpeg1^KO^ cells. Data represent mean and SEM of n = 4 independent experiments.

Unexpectedly, in cells treated with Z-FA-FMK to inhibit shedding, we saw no difference in the efficiency of saporin escape from phagosomes (Figure 5E, S3B,E). We were also not able to inhibit escape by interfering with the AEP-mediated cleavage in the P2 loop (Figure S3C,E), γ-secretase-mediated removal of the C-term fragment (Figure S3F), or with the promiscuous cleavage in the EGF domain (Figure S4). Together, these data suggest that neither shedding nor the cleavages in the EGF and P2 domains are required for endocytic escape from phagosomes.

### Perforin-2-mediated endocytic is not abolished by the V-ATPase inhibitor

Since shedding was inhibited in cells treated with BafA1 (see Fig 4D), we went on to test whether phagosome acidification is required for perforin-2-mediated antigen delivery into the cytosol. Interestingly, *in vitro*, pre-pore to pore transition of perforin-2 has been achieved by incubating pre-pores at pH 4 (Ni *et al*, 2020; Pang *et al*, 2019; Jiao *et al*, 2021). However, cDC1s, which are the main cross-presenting cells (Hildner *et al*, 2008), and employ perforin-2 to deliver antigens into the cytosol (Rodríguez-Silvestre *et al*, 2023), were reported to maintain their phagosomes at near-neutral pH (Savina *et al*, 2009; Netting *et al*, 2025; Mantegazza *et al*, 2008). Therefore, the requirement of pH 4 for the formation of endocytic escape pores appeared inconsistent with the physiology of cDC1 phagosomes. Using pH-sensitive beads coated with pHrodo, we confirmed that phagosomal pH in both primary and Flt3L-derived cDC1s remains neutral, in contrast to cDC2 or pDCs (Figure S6). Importantly, the pHrodo experiments suggest that cDC1-like MutuDCs, in contrast to primary cDC1s, efficiently acidify their phagosomes (Figure S5). Nevertheless, when treated with BafA1 to interfere with acidification, MutuDCs were still able to import saporin from phagosomes into the cytosol (Figure 5F).

Importantly, while treatment of MutuDCs with BafA1 significantly reduced phagosome acidification, a small increase in pHrodo fluorescence was still observed over time (Figure S6). This was in contrast to complete alkalisation of phagosomes with NH_4_Cl, which abolished escape (Figure S3G). Thus, a pH gradient during phagosome maturation is likely relevant for perforin-2 activity. Nevertheless, saporin escape in BafA1-treated MutuDCs as well as in primary and Flt3L-derived cDC1s which maintain phagosomes at near neutral pH, suggest that perforin-2-mediated endocytic escape is compatible with weak acidification of phagosomes. Considering shedding requires low pH, these data further support the model where full-length perforin-2 is employed during cross-presentation.

### Membrane anchoring by CTT is required for endocytic escape

If full-length perforin-2 mediates endocytic escape, the pores in phagosomes must be formed in the cis conformation, *i.e.* by rotating the MACPF domain towards the membrane on which the pre-pore sits (see Figure 1A) (Yu *et al*, 2022; Jiao *et al*, 2021). According to this model, the individual MAPCF domains flip and propagate insertion around the perforin-2 ring in a clockwise fashion. Interestingly, during this propagation, the intermolecular interactions between the two neighbouring subunits are almost completely lost, as in addition to the MACPF flip, the P2 domain rotates along the axis roughly perpendicular to the membrane (Yu *et al*, 2022).

Importantly, in the available structures of the pre-pores, the CTT domain wraps behind the P2 domain of the neighbouring subunit (Figure 6A). This has two important implications. First, it suggests that the M599^ shedding site in the CTT does not appear accessible to proteases as it is concealed by the P2 domain (Figure 6B). Second, it suggests that the CTT-P2 interaction might in fact provide the majority of intermolecular interactions between the subunits, necessary to maintain integrity of the ring during the MACPF flip (Figure 6A). Thus, to establish, whether the CTT is critical for endocytic escape, we introduced two charged residues, R609D/T610E or V609R/T610R, in the CTT β-hairpin that form contacts with the neighbouring P2 (Figure 6C). These point mutations significantly decreased the efficiency of the endocytic escape, and, surprisingly, the complete deletion of the β-hairpin, V607-K620Del, abolished escape (Figure 6D, S7), pointing at the critical role of the CTT during formation of endocytic escape pores.

**Figure 6.**
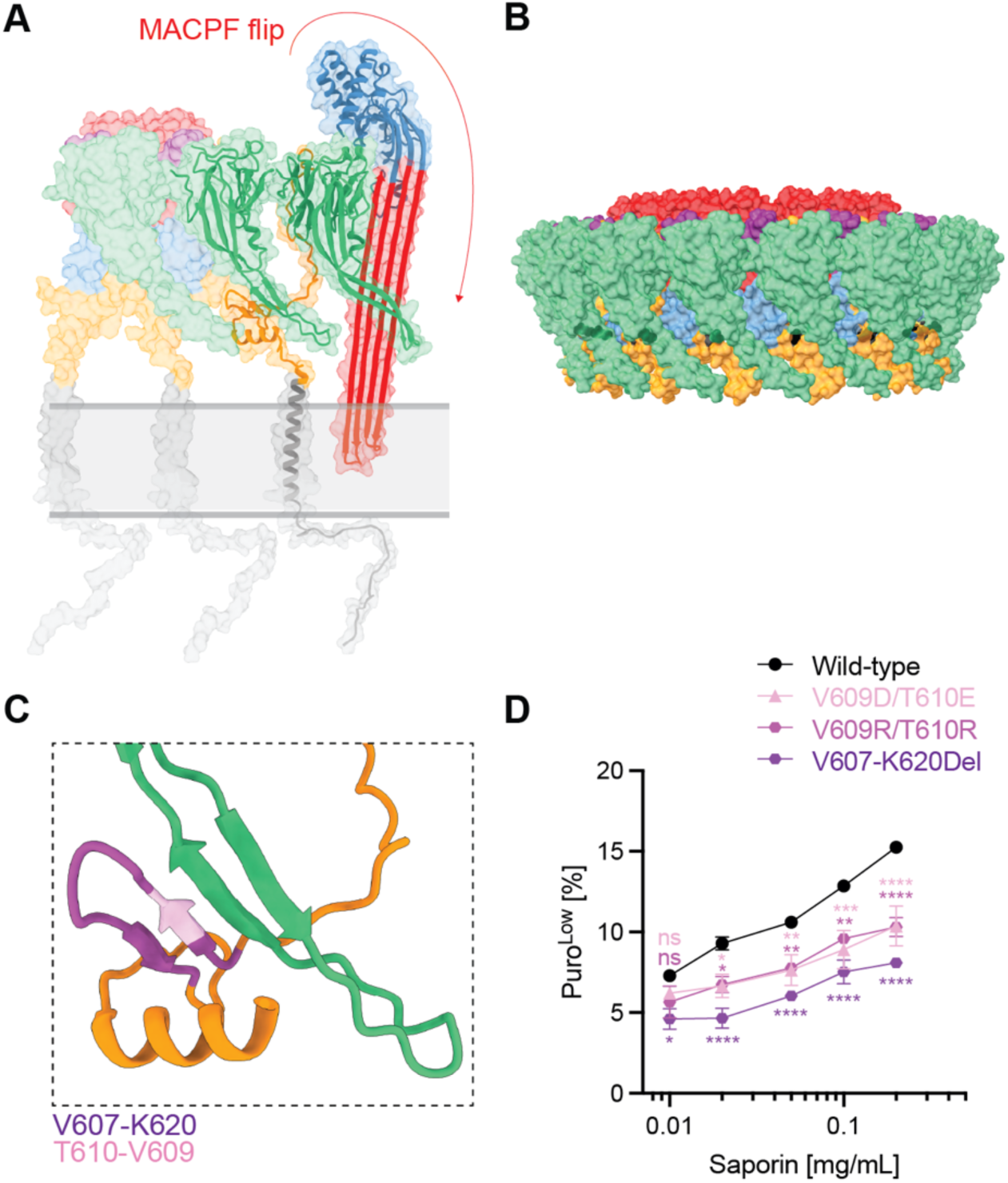
Anchoring via CTT domain is required for endocytic escape. **A** Model of the pre-pore to pore conformational transition of full-length perforin-2. **B** Shedding site M599^S600 is highlighted in black in the pre-pore structure of perforin-2 (pdb 6sb3). **C** CTT (orange) interaction with neighbouring P2 (green). Mutated residues are shown in pink and purple. **D** Mpeg1^KO^ MutuDCs were reconstituted with either WT Mpeg1 or with the indicated mutants and the bead-saporin phagosomal escape assay was performed. Data represent mean and SEM of n = 2 independent experiments.

## Discussion

Perforin-2 is the most ancient, yet least characterised, member of the MACPF family. It has been implicated in two fundamentally distinct processes: the killing of intravacuolar pathogens during innate immune responses and the translocation of antigens across endocytic membranes during cross-presentation and T cell priming. How these two perforin-2 functions are regulated remains unclear. Here, we set out to identify the mechanism that permits controlled release of phagocytic cargo without compromising integrity of intracellular compartments in dendritic cells. We demonstrate that anchoring of perforin-2 via the CTT and TMD to the phagosomal membrane is critical for escape.

The concept of regulating pore formation by proteolysis is well established for related pore-forming proteins. For example. perforin-1 requires the proteolytic removal of a C-terminal glycan that hinders oligomerisation (House *et al*, 2017; Uellner *et al*, 1997). The assembly of the complement membrane attack complex is regulated by a proteolytic cascade (Dunkelberger & Song, 2010) Similarly, members of the gasdermin family, which also form β-barrel pores, are activated through proteolytic activation by cytosolic caspases (Devant & Kagan, 2023). Our findings demonstrate that similar to other pore-forming proteins, perforin-2 also undergoes extensive proteolytic remodelling with at least five cleavage sites characterised through mass spectrometry and immunoblotting. These include loss of the N-terminal helix, AEP-mediated cleavages in the EGF and P2 domains that result in three disulfide-linked fragments, as well as shedding of the ectodomain and γ-secretase-mediated release of the TMD from the membrane. Importantly, in cDC1-like cell line MutuDC, proteolytically cleaved perforin-2 appears to be the major species.

Unexpectedly, we report that perforin-2-mediated endocytic escape was not compromised when we interfered with its proteolytic maturation. Instead, our data point at a critical role of membrane anchoring by the CTT and TMD during formation of pores in the endocytic compartments supporting the model where full-length perforin-2 mediates endocytic escape. The requirement of the TMD during pore formation is also consistent with the observation that the P2 domain does not bind mammalian lipids efficiently (Pang *et al*, 2019) and, therefore, it is unlikely that ectodomain alone can insert into self-membranes. According to the model we propose, it is the proteolytic release of the ectodomain, rather than TMD-mediated anchoring, that serves to protect the endogenous membranes once phagosomes acidify (see Figure S8 for a schematic). Importantly, since ectodomain shedding occurs in low pH compartments, it would explain how the dendritic cell cytosol is protected from hydrolytic enzymes present in acidified perforin-2-containing phagosomes or endolysosomes. We speculate that rapid acidification of phagosomes in other *Mpeg1-*expressing cell types, such as macrophages, might favour P2 domain flipping revealing the shedding site and formation of proteolytically-released trans pores in intravacuolar pathogens.

The mechanism of endocytic escape, where full length perforin-2 forms cis pores in weakly acidified compartments to facilitate antigen escape explains several discrepancies in the proteolytic release model. (1) The proteolytically released ectodomain behaves as a soluble rather than a membrane-inserted protein in low pH compartments. (2) The proton gradient across the phagosomal membrane is maintained during perforin-2-mediated endocytic escape. (3) cDC1s do not acidy their phagosomes, yet they employ perforin-2 during cross-presentation. (4) A pre-pore lacking a TMD is unlikely to be able to associate with phagocytic membranes since the tip of the P2 domain shows preferential binding to bacterial rather than mammalian lipids. (5) The P2 domain is located in front of the CTT domain in the pre-pore conformation protecting the shedding site from proteases and making it implausible that shedding could occur prior to cis pore formation.

If ectodomain shedding by lysosomal enzymes is not the trigger for pore formation, what drives the formation of the cis pores? *In vitro*, perforin-2 pores were obtained by incubating pre-pores formed by perforin-2 ectodomain at pH 4 (Jiao *et al*., 2022). It is possible that these pores were formed in trans (by flipping of the P2 domain) rather than in cis (by flipping of the MACPF domain) and represent pores formed in acidic lysosomes to attack intravacuolar pathogens. Indeed, trans pores bridging two liposomes have been observed in cryo-electron microscopy data (Pang *et al*, 2019). Furthermore, it is still possible that in the early endocytic compartments, the mild drop of pH to ∼6 is sufficient to drive formation of the cis pores in the context of full-length perforin-2. Indeed, our data showed that while treatment with V-ATPase inhibitor BafA1 does not abolish endocytic escape, complete neutralisation of endocytic compartments with NH_4_Cl does.

In summary, we demonstrate that full length perforin-2 mediates antigen delivery into the cytosol of dendritic cells. We also propose that endocytic escape occurs via cis pores forming in weakly acidified endocytic compartments, and that pH drop favours formation and proteolytic release of trans pores which attack pathogens. Thus, the TMD enables context-dependent functions of perforin-2 and repurposing the same effector for fundamentally different tasks in different cell types. This dual function might reflect the ancestral origin of perforin-2, from which complement and perforin-1 later diverged as specialised effectors for microbial killing and delivery of effectors across mammalian membranes.

## Materials and Methods

### Animals

All mice were bred/maintained in pathogen-free conditions by the Medical Research Council ARES facility. Experiments were approved by the LMB Animal Welfare and Ethical Review Body, and the UK Home Office approved experiments. C57BL/6 (Rodríguez-Silvestre *et al*, 2023) (female or male) and Mpeg1^−/-^ (Rodríguez-Silvestre *et al*, 2023) mice used were 8 to 13 weeks old when culled for tissue collection.

### Antibodies and reagents

All antibodies and reagents used in this study are listed in Table S2.

### Cell lines and cell culture

MutuDCs were a kind gift from Hans Actha-Orbea. The cells were maintained in IMDM supplemented with 8% heat-inactivated FBS, 10 mM HEPES, 500 μM 2-mercaptoethanol, 1X GlutaMAX, penicillin (100 units/mL) and streptomycin (100 μg/mL) at 37°C and 5% CO_2_. HEK-293Ts were maintained in DMEM supplemented with 10% heat-inactivated FBS and 1X GlutaMAX at 37°C and 5% CO_2_.

### Generation of Flt3L-BMDCs

Bone marrow was recovered from femurs and tibiae of from 8–13-week-old male or female C57BL/6J mice and resuspended in in RPMI-1640 supplemented 10% heat inactivated FBS, 10 mM HEPES, 500 μM 2-mercaptoethanol, 1X sodium pyruvate, 1X non-essential amino acids, penicillin (100 units/mL) and streptomycin (100 μg/mL). For Flt3-L cultures, 15×10^6^ cells were seeded in a non-treated 10 cm dish and supplemented with 5 ng/mL GM-CSF and 200 ng/mL Flt3-L (Day 0). Cells were differentiated for 10 days with the addition of 10 mL of fresh media with 5 ng/mL GM-CSF and 200 ng/mL Flt3-L on Day 5 and 5 mL of fresh media containing 20 ng/mL GM-CSF on Day 7.

### Isolation of splenic DCs

Spleens were perfused with a solution of RPMI-1640, 0.1 mg/mL Liberase-TL and 0.1 mg/mL DNAse I, minced and digested for 25 min at 37_°_C. Heat-inactivated FBS was added at 10% (v/v) to stop the digestion, before mashing tissues through a 70 μm filter. Red blood cells were lysed with Hybri-Max for 3 min at room temperature. Dendritic cells were isolated using a Miltenyi Pan-DC Isolation Kit (negative selection) according to the manufacturer’s instructions.

### Generation of Mpeg1 mutants

pHR-mMpeg1-IRES-mScarlet-V609D/T610E, pHR-mMpeg1-IRES-mScarlet-V609R/T610R and pHR-mMPEG1-IRES-mScarlet-V607-K620del Mpeg1 variants were generated by cloning custom GeneBlock DNA fragments (IDT, see Table S2 for sequences) into pHR-MPEG1-IRES-mScarlet vector (Rodríguez-Silvestre *et al*, 2023). Vectors and gene fragments were digested using *Sbf*I and *Blp*I restriction for 2 h at 37°C. Digested vectors were gel-purified from a 1% agarose gel using the QIAquick Gel Extraction Kit, and digested gene fragments were purified using the QIAquick PCR Purification Kit. Ligations were performed using T4 DNA ligase according to the manufacturer’s instructions. Transformations were performed using Top10 chemically competent E. coli according to the manufacturer’s instructions.

### Generation of sgRNA expression plasmids

sgRNAs were designed using the VBC score tool (https://www.vbc-score.org/) by choosing the highest ranked sgRNA for each gene (Michlits *et al*, 2020). The chosen sgRNAs target exon 1, exon 8 and exon 6 of Aph1c, Ncstn and Psen2, respectively. To generate sgRNA expression plasmids, lenti-sgRNA-hygro (Table S2) was digested with BsmBI, dephosphorylated using Quick CIP according to the manufacturer’s instructions and purified using the QIAquick PCR purification kit. The sgRNA oligos were purchased as IDT RxnReady Primer Pools (Table S2). The oligos were phosphorylated using T4 Polynucleotide Kinase according to the manufacturer’s instructions, heated to 95°C for 5 min to ensure hybridisation, and inserted into lenti-sgRNA-hygro plasmid.

### Lentiviral transduction of MutuDCs

For lentiviral production, 3×10^6^ HEK-293T cells were seeded into a 10 cm dish. The next day, the cells were transfected with a mixture of 45 μL TransIT-LTI, 7 μg of the expression vector with the gene of interest along with 7 μg psPAX2 and 0.7 μg pMD2.g in 1 mL of OptiMEM. The medium was replaced 18 h post transfection with fresh medium supplemented with 1% BSA. After 48 h post transfection, viral supernatant was collected, passed through a 0.45 μm-syringe filter and 10X concentrated using Amicon Ultracentrifugal filters with a 100,000 MW cut-off. Aliquots were stored at -80°C. For transduction, 2.5×10^6^ MutuDCs were seeded in a 6 cm dish in 3 mL of media. After letting the cells adhere for at least 2 h, 1 mL of 10X concentrated virus was added in the presence of 6 μg/mL polybrene. The next day, the virus was removed and replaced with fresh culture medium. Two days post-transduction, cells were selected with 200 μg/mL hygromycin for one week. *Mpeg1* mutants were transduced into Mpeg1^KO^ MutuDCs and sgRNA guides for creating γ-secretase deficient cells were transduced into Cas9-expressing MutuDCs (Rodríguez-Silvestre *et al*, 2023). To test the knockout efficiency, genomic DNA was isolated and regions around the cut sites were amplified by PCR (Table S2). The amplicons were sequenced and analysed using TIDE (http://shinyapps.datacurators.nl/tide/).

### Preparation of ovalbumin-coated beads (Ova beads) for the phagoFACS assay

Amino-modified microspheres (Table S2) with a diameter of 3 μm were washed twice in PBS and preactivated with 8% (vol/vol) glutaraldehyde for 4 h at room temperature. Preactivated beads were washed once in PBS and then incubated overnight at 4°C with ovalbumin (Table S2) at a concentration of 0.5 mg/mL in PBS. The next day, beads were quenched in 0.4 M glycine in PBS for 30 min, washed twice in PBS and used immediately for phagocytosis.

### Preparation of saporin-coated beads

Saporin (Table S2) and BSA (Table S2) were conjugated to Ova beads through disulfide bonds. For efficient conjugation, free sulfhydryl groups were introduced by reacting 0.6 mL of 2.5 mg/mL saporin or BSA with 12 μL of 2 mg/mL Traut’s reagent in PBS containing 2 mM EDTA for 60 min at room temperature. Excess reagent was removed using a Zeba spin desalting column (Table S2) equilibrated with PBS containing 2 mM EDTA. Ova beads were resuspended in PBS containing 2 mM EDTA and reacted with 1 mM SPDP for 30 min at room temperature. The beads were then washed twice in PBS containing 2 mM EDTA and incubated with different mixtures of cysteine-modified saporin and BSA overnight at room temperature. Saporin was titrated across the mixtures (1 mg/mL to 0 mg/mL) and substituted with BSA so that the total protein concentration in each mixture remained constant at 1 mg/mL. The next day, beads were washed three times in PBS and immediately used for phagocytosis.

### Preparation of pHrodo-coated beads

Amino-modified microspheres with a diameter of 3 μm were washed twice in PBS, resuspended in 100 mM sodium bicarbonate (pH 8.5) and reacted with 0.2 mM pHrodo iFL Red Ester dye (Table S2) for 1 h at room temperature. The beads were then washed once in PBS, and any reactive ester moieties were quenched by incubating the beads in 1X TBS for 10 min. After an additional two washes in PBS, the beads were coated with 1 mg/mL ovalbumin (Table S2) by passive absorption for 1 hour at room temperature. The beads were then washed three times in PBS and immediately used for phagocytosis.

### PhagoFACS assay

MutuDCs were collected, washed once in PBS and resuspended in ice-cold internalisation medium (CO_2_-independent medium containing 1X GlutaMAX) to a density of 20×10^6^ cells/mL. Ova beads were added at a 10:1 ratio of beads: cells and incubated for 25 min at 16°C followed by a 5-minute incubation at 37°C to allow phagocytic binding and internalisation of beads. To remove non-internalised beads, cells were first washed twice with 10 mL ice-cold PBS at 100 x g for 4 min at 4°C and then resuspended in 1 mL PBS, applied to a 5 mL FBS cushion and centrifuged at 150 x g for 4 min at 4 °C. The cell pellet was then resuspended to 20×10^6^ cells/mL in cell culture medium and divided into different time points comprising 5×10^6^ cells each. For drug treatments, the culture medium was supplemented with 0.5 μM bafilomycin A1, 0.1 μM brefeldin A, 10 μM DAPT or 10 μM Z-FA-FMK. The chase was performed at 37°C for different periods of time and stopped by adding ice-cold PBS. Non-internalised beads were stained on ice with a goat αOvalbumin antibody (1:100) (see Table S2 for all antibodies) for 30 min followed by an αGoat-AF488 antibody (1:200) for 30 min. Cells from each time point were resuspended in 0.5 mL homogenization buffer (250 mM sucrose, 3 mM imidazole, 2 mM DTT, 2 mM PMSF and 1X protease inhibitor cocktail, pH 7.4) and passed 25 times through a 22-G needle. Intact cells and debris were pelleted by centrifugation at 150g for 4 min and the phagosome-containing post-nuclear supernatants transferred to a V-bottom 96-well plate. The enriched phagosomes were washed with PBS containing 1% (vol/vol) BSA and stained with different primary antibodies overnight at 4°C. The next day, the samples were incubated with appropriate secondary antibodies for 45 min on ice. Phagosomes were analysed by flow cytometry using a CytoFlex flow cytometer. The data were analysed using FlowJo software.

### Bead saporin-puromycin phagosomal escape assay

The bead saporin-puromycin was performed in cell culture medium lacking β-mercaptoethanol (βM-free) to avoid reduction of the conjugating disulfide bond on saporin beads. MutuDCs were collected, washed once in PBS and seeded in a 96-well U-bottom plate with 5×10^5^ cells per well in 100 µL βM-free cell culture medium. Saporin beads were added at a beads:cells ratio of 10:1 in 50 µL βM-free medium. For drug treated cells, the βM-free medium was supplemented with a final concentration of 0.5 µM bafilomycin A1, 10 μM DAPT or 10 μM Z-FA-FMK. Cells were incubated with beads at 37 °C for the indicated times. Then, puromycin was added at 10 µg/mL for 30 min, and cells were subsequently washed in ice-cold PBS. Non-internalised beads were stained on ice with a rabbit αOvalbumin antibody (1:200) for 30 min followed by an αRabbit-AF455 antibody (1:1000) for 30 min. After labelling dead cells with ViaKrome 808 (1:1000) for 10 min on ice, cells were fixed and permeabilised using the BD Cytofix/Cytoperm and Perm/Wash buffers. Puromycin incorporation was determined by staining with an αPuromycin-AF647 antibody (1:200) in Perm/Wash buffer for 45 min on ice. Cells were analysed by flow cytometry using a CytoFlex flow cytometer. The data were analysed using FlowJo software.

### pHrodo assay

For pHrodo assays with MutuDCs and BMDCs, the pulse with pHrodo beads was performed as described in Section 2.5.1. For pHrodo assays with splenic DCs, the pulse was adjusted to 15 min for 37°C to allow comparison to a previous study (245). The chase was performed at 37°C for different periods of time and stopped by adding ice-cold PBS. For drug treated cells, the medium was supplemented with a final concentration of 0.5 µM bafilomycin A1 or 5 μM DPI. Non-internalised beads and dead cells were labelled in PBS containing 1% (vol/vol) BSA with a rabbit anti-OVA antibody for 30 min on ice followed by a staining with donkey anti-rabbit-AF647 and ViaKrome 808 for 30 min on ice. For pHrodo assays with Flt3L-BMDCs and splenic DCs, non-specific antibody binding was blocked using αCD16/CD32 prior to staining. The staining also included antibodies against surface markers. Stained cells were immediately analysed on a CytoFlex flow cytometer using a chilled sample stage. The data were analysed using FlowJo software.

### Western blotting

For western blotting of drug treated cells, MutuDCs were seeded into a 6-well dish at 2×10^6^ cells/well in 2 mL of cell culture media. The next day, cells were incubated with inhibitors as indicated in the relevant figure legends. Drug-treated, or untreated cells, were pelleted and washed once in ice-cold PBS. Pellets were lysed in RIPA buffer supplemented with 1X protease inhibitor cocktail for 20 min at 4°C while shaking at 800 rpm. Insoluble material was pelleted by centrifugation at 20,000g for 10 min. Supernatants were mixed with NuPAGE LDS Sample Buffer and heated at 70 °C for 10 min in the presence or absence of 1X Bolt reducing agent. Samples were run in MOPS buffer on NuPAGE 4–12% Bis-Tris gels at 150V for 60 min and transferred onto nitrocellulose membrane using an iBlot1 system. Loading was controlled by Ponceau staining. Membranes were then blocked by incubation with blocking buffer (PBS containing 5% milk 0.1% Tween-20) for 1 hour at room temperature. Primary antibodies were diluted in blocking buffer and incubated at 4°C overnight. After washing three times with PBS containing 0.1% Tween-20, membranes were incubated with HRP-conjugated secondary antibodies diluted in blocking buffer for 1 hour at room temperature. Membranes were again washed three times with PBS containing 0.1% Tween-20 and then incubated with ECL reagent for 10 min. The resulting signal was detected using X-ray films with varying exposure times.

### Identification of cleavage sites from mass-spectrometry data

Proteomic data from (Rodríguez-Silvestre *et al*, 2023) was used for identification of cleavage sites in perforin-2. Three control samples from the experiment with small molecules and three NTC control samples from the experiment with AEP^KO^ lines were used for analysis. Non-tryptic peptides were filtered for those with LFQ intensity > 25 in each of the six replicates.

### Identification of the predicted cleavage sites

Mouse perforin-2 sequence was used for analysis using The Eukaryotic Linear Motif resource, 2024 update (version 2.6.1) (https://elm.eu.org/) (Kumar *et al*, 2023). Mouse perforin-2 sequence was used for analysis.

### Structural analyses

Molecular graphics and analyses performed with UCSF ChimeraX (version 1.8) (Meng *et al*, 2023). To model a structure of the full length pre-pore monomer from pdb 6SB3 (Ni *et al*, 2020) was aligned with Alphafold model of perforin-2, AF-A1L314-F1 (Jumper *et al*, 2021)using the matchmaker function, and the TMD was manually rotated to place into the membrane on which the pre-pore sits (residues 643-713 are shown from the Alphafold structure). Full length trans pore was modelled in a similar way using pdb 8AD1 (Yu *et al*, 2022) (residues 600-713 are shown from the AlphaFold structure).

To model the pre-pore to cis pore transition, we used three subunits from the full-length pre-pore model created above. P2 (374-577) domains of the rotating subunit from pre-pore and pore were aligned using matchmaker. In the rotating subunit residues 1-583 are derived from 8a1d, 584-642 from 6sb3, and 643-713 from the Alphafold structure.

### Statistical analysis

Details of the statistical analysis are provided in Figure Legends. Plots were generated using GraphPad Prism version 10.6.0 for Mac (GraphPad Software, La Jolla California USA).

## Acknowledgements

We would like to thank Dr. Felix Randow and Dr. Leo James for advice and all the members of the Kozik lab for helpful discussions and feedback. We are grateful to the Flow Cytometry Core, Genotyping, and Biological Services (Ares) Facilities at MRC Laboratory of Molecular Biology for help and technical assistance. We would also like to thank Pablo Rodiguez-Silvestre for help with experiments involving primary DCs. This project was supported by the Medical Research Council, as part of United Kingdom Research and Innovation (also known as UK Research and Innovation) (MC_UP_1201/26).

## Author contributions

ML and PK conceived the study. ML designed and carried out the majority of experiments with the exceptions detailed below. RC performed some of the western blots. PK performed the computational analysis of the mass spectrometry data and the structural analysis. ML, RC, and PK wrote and edited the manuscript.

