## Supplementary Figures for "Anchoring of perforin-2 via the transmembrane domain is required for endocytic escape in cross-presenting dendritic cells"

### List of Supplementary materials

Table S1. Cleavage sites and candidate proteases predicted by the ELM resource (Excel).

Table S2. Reagents and tools

Supplementary Figures (pdf).

**Supplementary Figures**

**Figure S1**

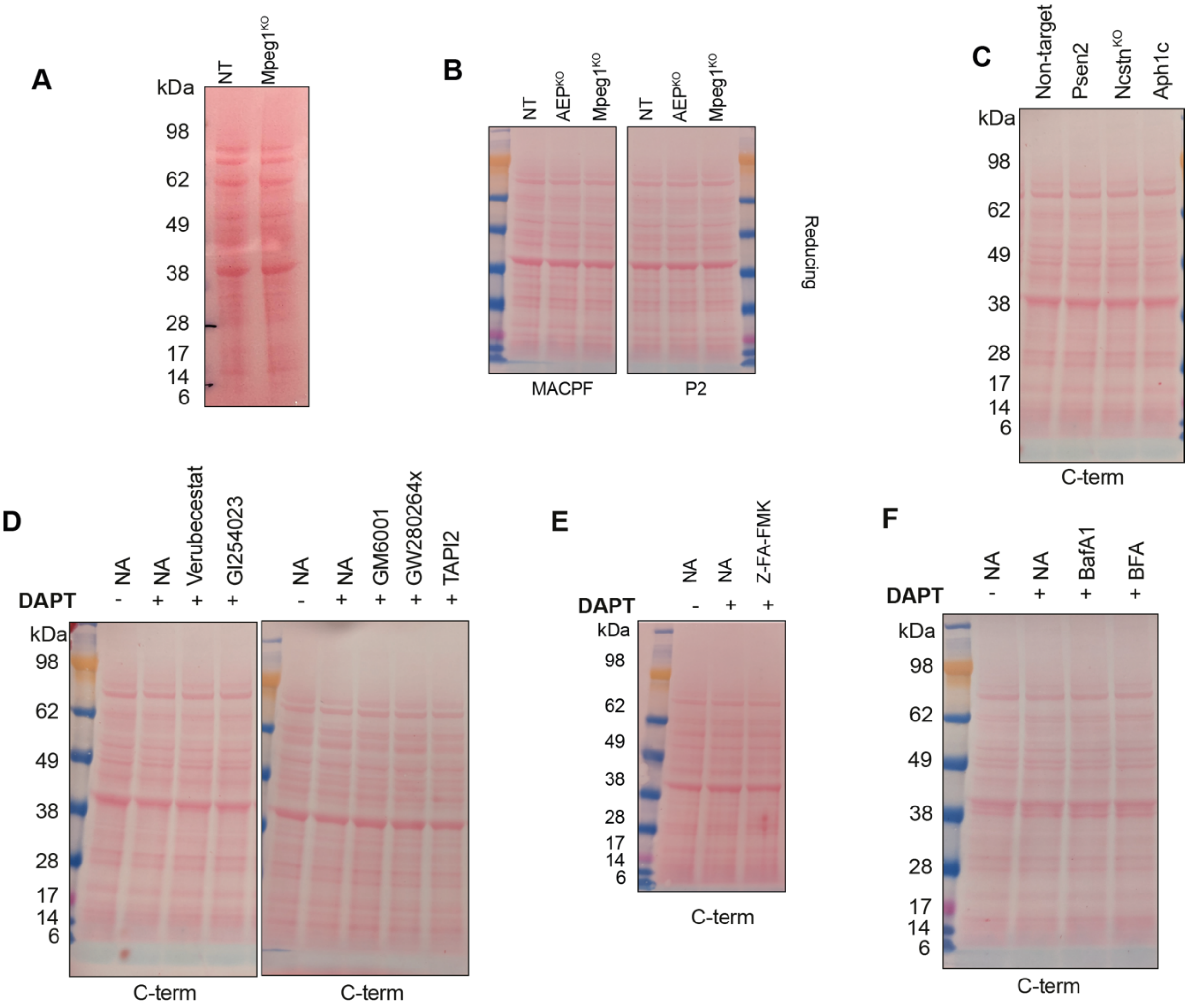


**Figure S1. Ponceau-stained blots for Western Blots included in the manuscript.**

**A** Loading control for Figure 2B
**B** Loading control for Figure 2C
**C** Loading control for Figure 3B
**D** Loading control for Figure 4C
**E** Loading control for Figure 4E
**F** Loading control for Figure 4D

**Figure S2**

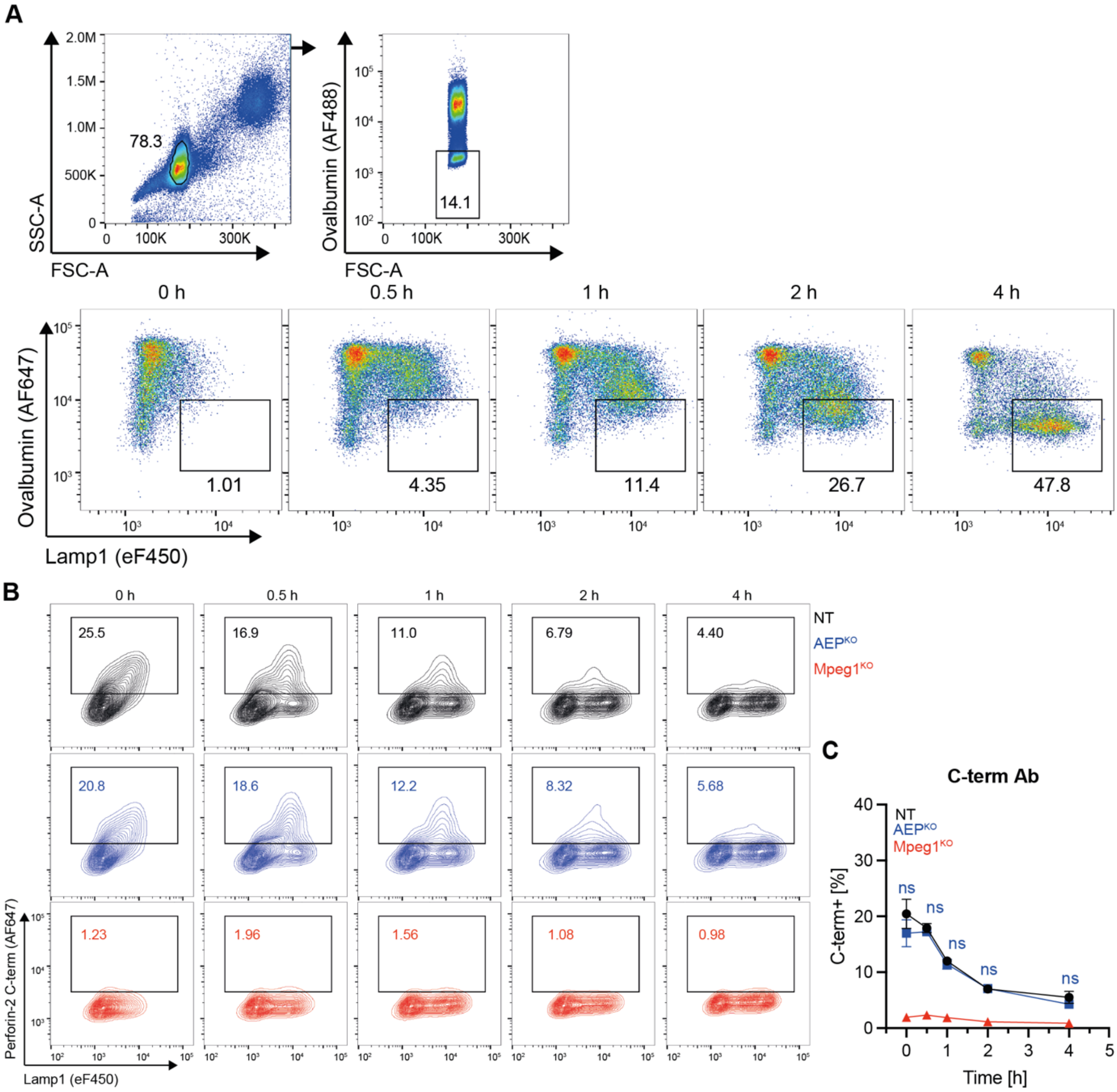


**Figure S2. Gating strategy for phagoFACS experiments and analysis of perforin-2 recruitment with the C-term antibody.**

**A** Gating strategy for NT MutuDCs were pulsed with OVA beads for 25 min at 16°C followed by 5 min at 37°C and either immediately placed on ice (0 h) or chased for the indicated time. Isolated phagosomes were stained with antibodies against ovalbumin and Lamp1. The gate indicates mature phagosomes. Data are representative of n =5 independent experiments.

**B** The indicated MutuDC lines were pulsed with Ova beads for 25 min at 16°C followed by 5 min at 37°C, washed and either immediately placed on ice (0 h) or chased for the indicated time. Isolated phagosomes were stained with αLamp1 and with αC-term antibody.

**C** Quantification of perforin-2+ phagosomes based on the gating shown in **B**. Data represent mean and SEM of n = 3 independent experiments.

**Figure S3**

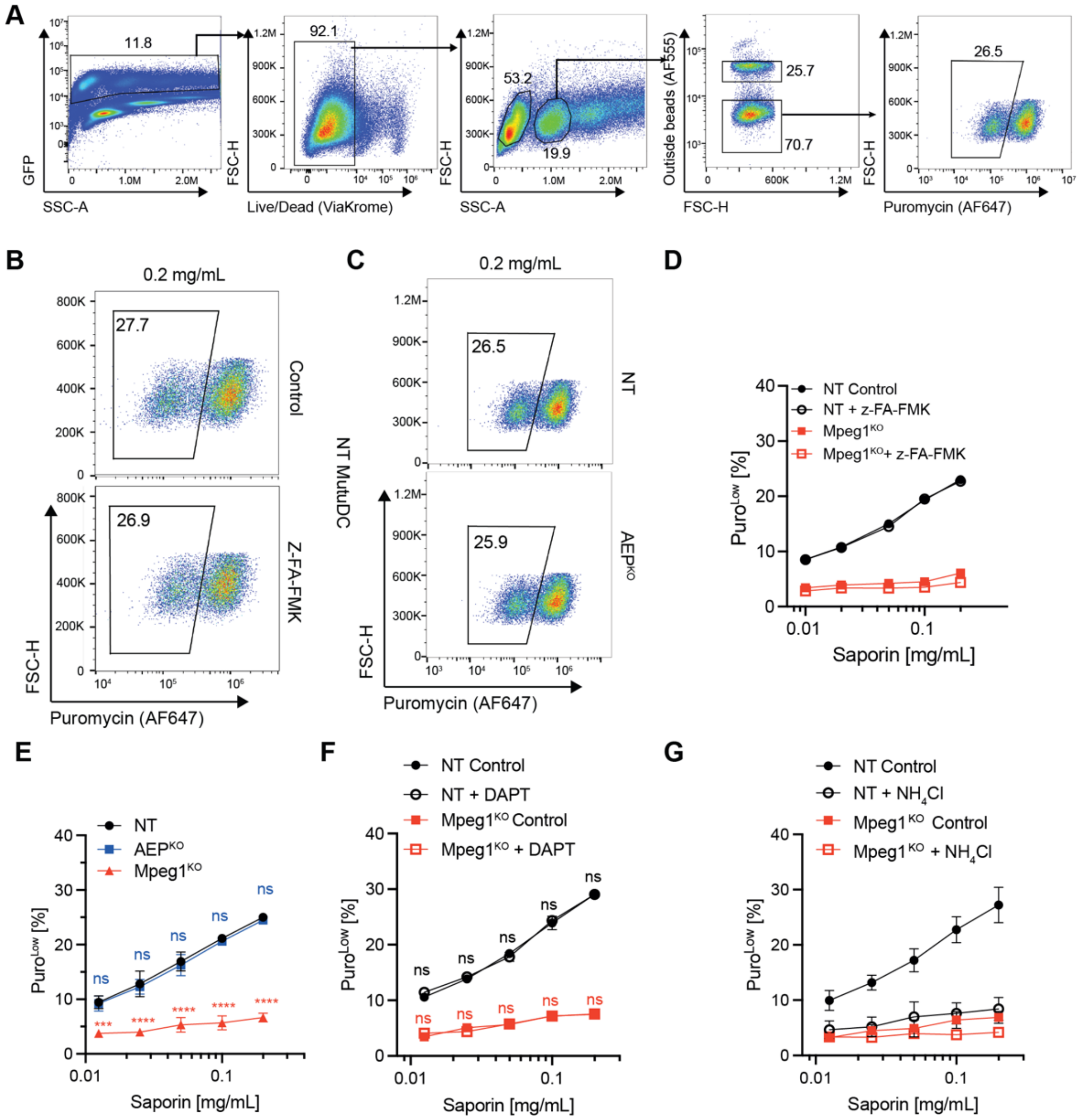

**Figure S3. Saporin-puromycin phagosomal escape assay.**

**A** Gating strategy for the saporin-puromycin assay to monitor antigen escape from phagosomes.

**B** Representative flow plot gating for PuroLow population for the saporin phagosomal escape assay in NT MutuDCs with or without pre-treatment with Z-FA-FMK for 1h.

**C** Representative flow plots gating for PuroLow population for the bead saporin-puromycin assay in NT or AEP $^{KO}$ MutuDCs.

**D** NT or Mpeg1$^{KO}$ MutuDCs were incubated with saporin beads for 5 h followed by an incubation with puromycin for 30 min and pre-incubated for 16h with Z-FA-FMK. Cells were then placed on ice and outside beads labelled with an αOvalbumin antibody. After fixation and permeabilisation, levels of puromycin incorporation were detected with an αPuromycin antibody. Quantification of cells in translation arrest (PuroLow).

**E** Quantification of PuroLow cells in bead-saporin escape assay in NT, $AEP^{KO}$, and Mpeg1 $^{KO}$ MutuDCs. n = 3 independent experiments.
**F** Quantification of PuroLow cells in bead-saporin escape assay in NT and $Mpeg1^{KO}$ MutuDCs with and without treatment with DAPT. n = 3 independent experiments.
**G** Quantification of PuroLow cells in bead-saporin escape assay in NT and $Mpeg1^{KO}$ with and without treatment with $NH_4Cl$ MutuDCs. n = 3 independent experiments.

### Figure S4

**A**

| | |
|---|---|
| z-FA-FMK | Cysteine proteases |
| Leupeptin | Proteases inhibitor |
| E64D | Cysteine proteases-Cathepsin and Calpain |
| CA-074 | Cathepsin B inhibitor |
| CA-074-Me | Cathepsin B and L inhibitor |

**B**

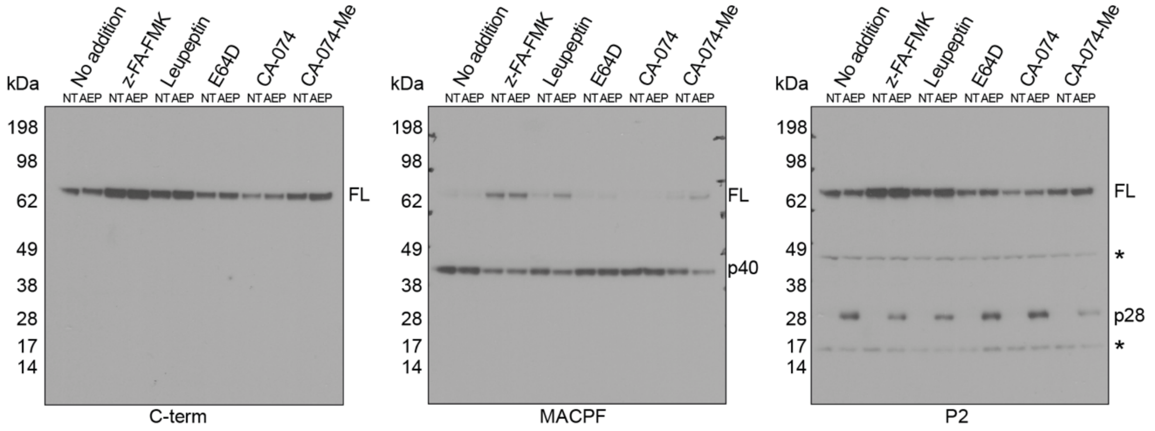


**C**

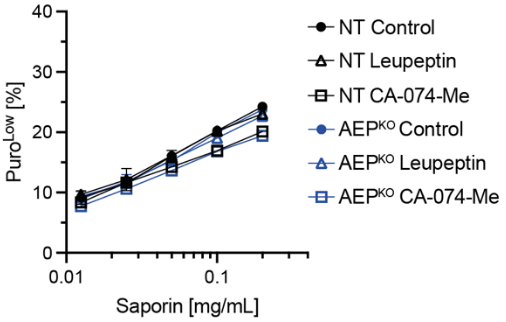


**Figure S4. Promiscuous cleavage in the EGF domain is not required for saporin escape.**

**A** Small-molecule inhibitors and their targets.

**B** NT or AEP$^{KO}$ MutuDCs were either left untreated or treated with Z-FA-FMK (10 µM), Leupeptin (100 µM), E64D (10 µM), CA-074 (50 µM) or CA-0740-Me (50 µM) for 6 hours and lysates analysed by immunoblot under reducing conditions using the αC-terminal tail, αMACPF or αP2 antibody. Leupeptin and CA-074-Me have a synergistic effect with AEP depletion, suggesting that cleavage in the EGF linker is promiscuous and can be executed by either AEP or cathepsins. n = 2 independent experiments.

**C** Quantification of PuroLow cells in bead-saporin escape assay in NT and AEP$^{KO}$ MutuDCs with and without treatment with the leupeptin or CA-074-Me. These data support the notion that cleavage in the EGF domain is not required for endocytic escape. Data represent mean and SEM of n =2 independent experiments.

**Figure S5**

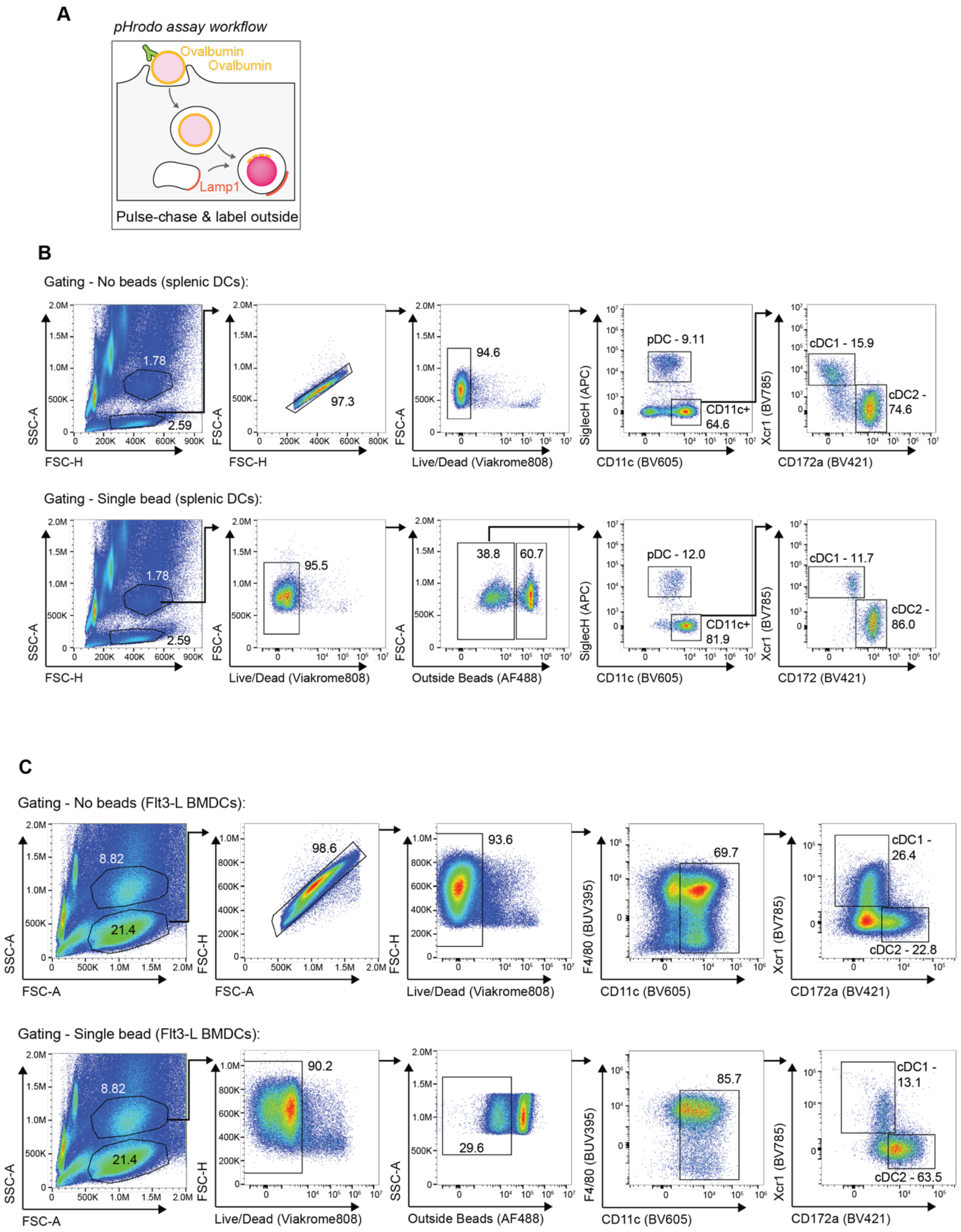
A
pHrodo assay workflow
Ovalbumin
Ovalbumin
Lamp1
Pulse-chase & label outside
B
Gating - No beads (splenic DCs):
1.78
2.59
97.3
94.6
pDC - 9.11
CD11c+
64.6
cDC1 - 15.9
cDC2 -
74.6
SSC-A
FSC-H
FSC-A
Live/Dead (Viakrome808)
SiglecH (APC)
CD11c (BV605)
Xcr1 (BV785)
CD172a (BV421)
Gating - Single bead (splenic DCs):
1.78
2.59
95.5
38.8
60.7
pDC - 12.0
CD11c+
81.9
cDC1 - 11.7
cDC2 -
86.0
Outside Beads (AF488)
CD172 (BV421)
C
Gating - No beads (Flt3-L BMDCs):
8.82
21.4
98.6
93.6
69.7
cDC1 -
26.4
cDC2 - 22.8
SSC-A
FSC-A
FSC-H
Live/Dead (Viakrome808)
F4/80 (BUV395)
CD11c (BV605)
Xcr1 (BV785)
CD172a (BV421)
Gating - Single bead (Flt3-L BMDCs):
8.82
21.4
90.2
29.6
85.7
cDC1 -
13.1
cDC2 - 63.5
Outside Beads (AF488)

**Figure S5 (continued)**

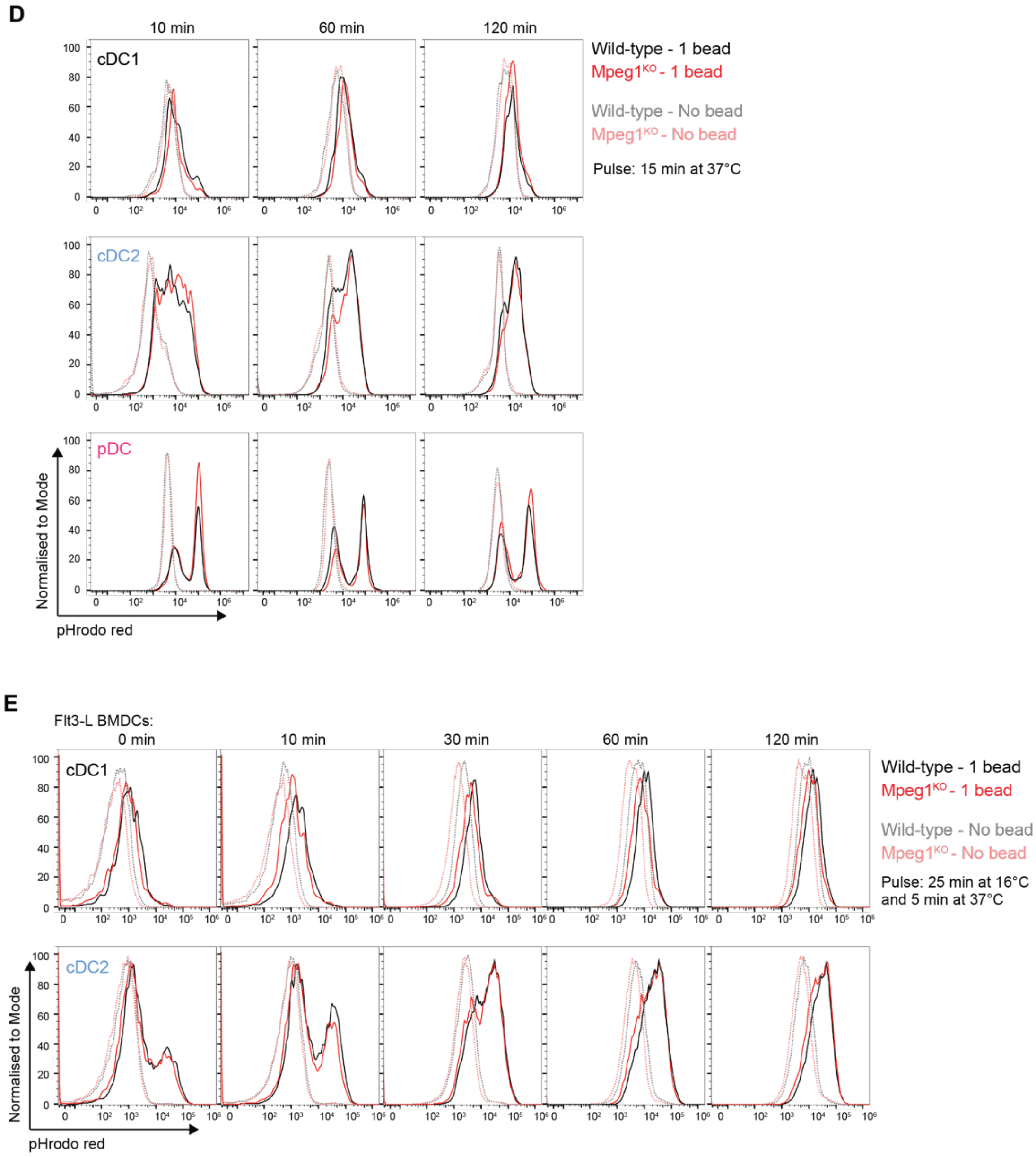


### Figure S5. pHrodo assay in primary and Flt3L-derived DCs.

**A** Schematic representation of the pHrodo bead assay.

**B** Gating strategy for the pHrodo bead assay in primary splenic DCs.

**C** Gating strategy for the pHrodo bead assay in primary Flt3L-bone marrow derived dendritic cells (Flt3-BMDCs)

**D** Wild-type or Mpeg1$^{KO}$ splenic DCs were pulsed with pHrodo beads for 15 min at 37°C and chased for the indicated time Data are representative for n = 2 independent experiments, respectively.

**E** Same as **D** for Flt3L-derived BMDCs.

### Figure S6

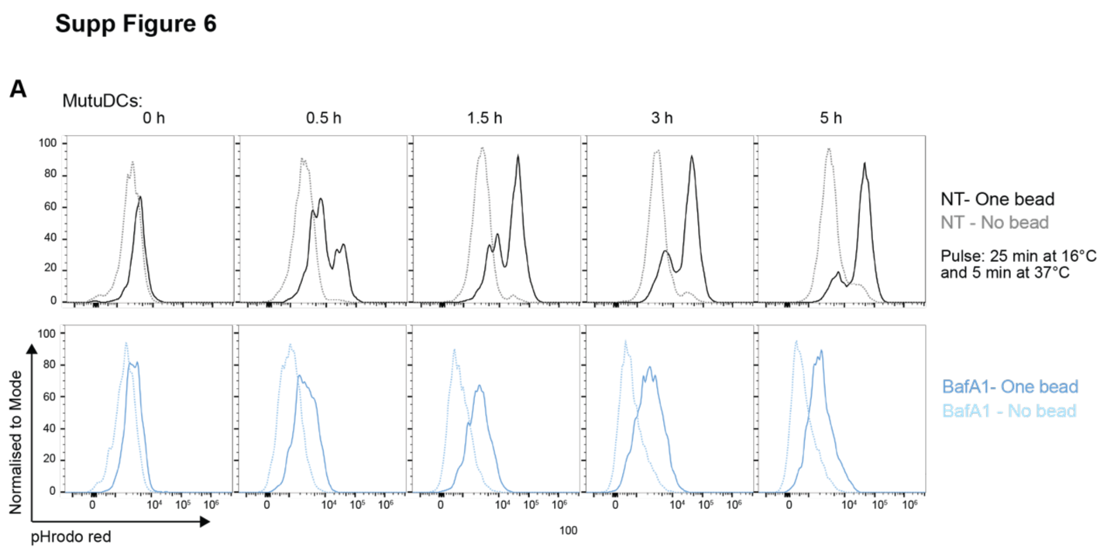


**Figure S6. pHrodo assay in control and BafA1-treated MutuDCs.**

**A** NT MutuDCs were subjected to the pHrodo bead assay as described in Sup Fig 6B in the presence or absence of 0.5 μM BafA1. Data representative of n = 5 independent experiments.

**Figure S7**

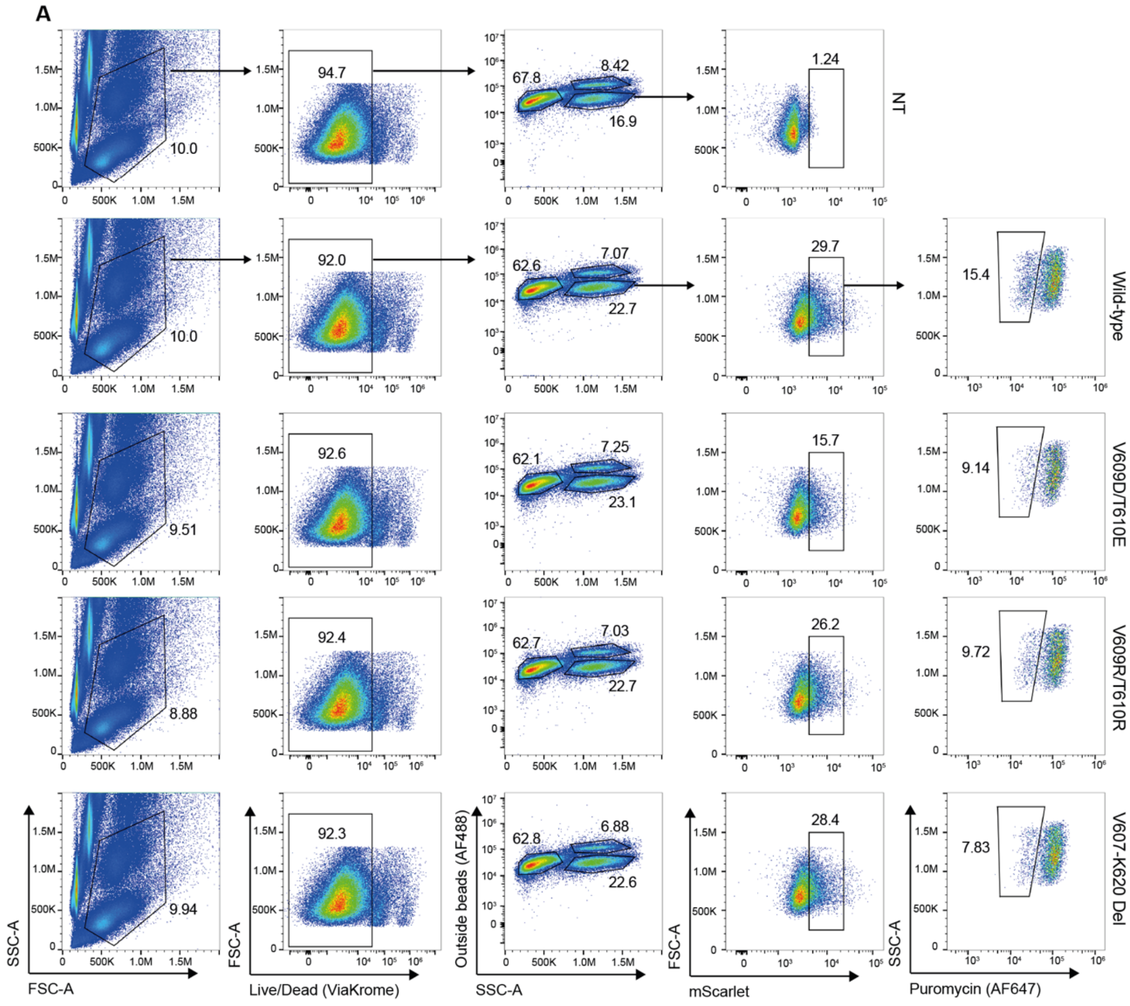


**Figure S7. Gating strategy for saporin assay with the Mpeg1 CCT mutants.**

**A** Gating strategy for the indicated MutuDC lines: NT (control), and Mpeg1$^{KO}$ MutuDC reconstituted with WT, V609D/T610E, V609R/T610R and V607-K620 Del perforin-2 mutants. The cells were incubated with saporin-beads for 5 h followed by an incubation with puromycin for 30 min. Cells were then placed on ice and outside beads labelled with an αOvalbumin antibody. After fixation and permeabilisation, levels of puromycin incorporation were detected with an αPuromycin antibody. mScarlet levels were used to normalise for transduction efficiency. Data represent the mean and SEM of n = 2 independent experiments.

### Figure S8

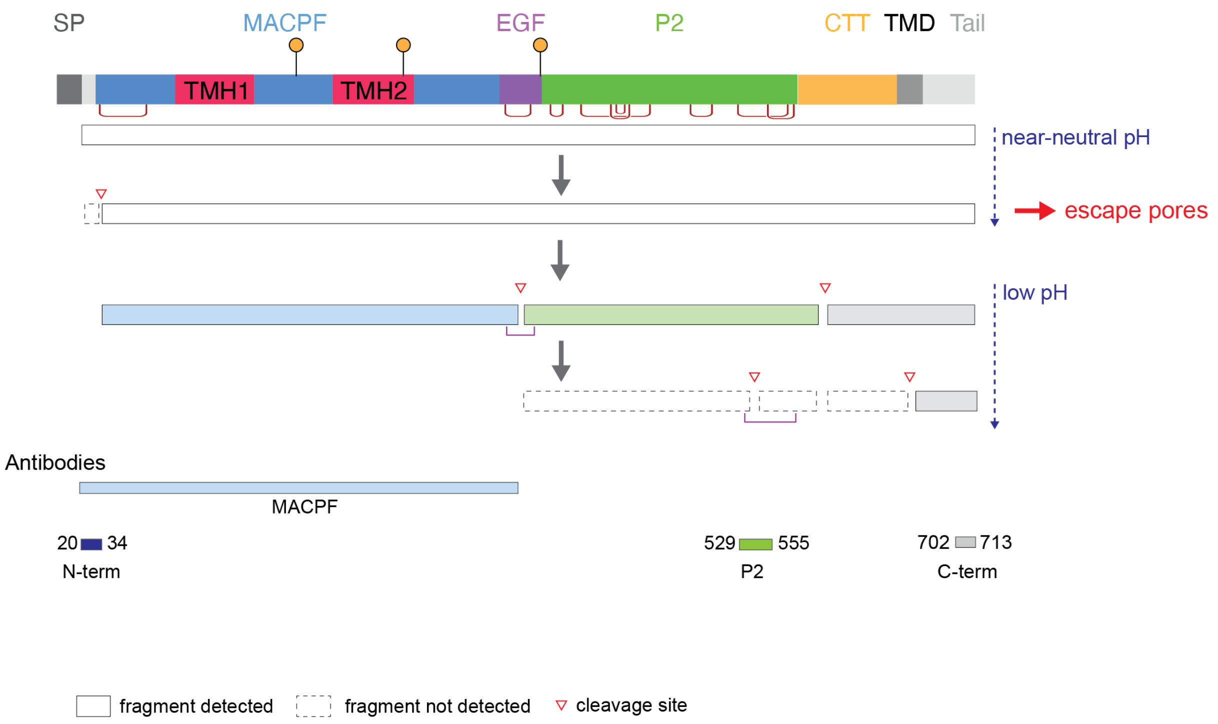


**Figure S8. Steps involved in perforin-2 processing.**
